# Behavioral control by depolarized and hyperpolarized states of an integrating neuron by

**DOI:** 10.1101/2021.02.19.431690

**Authors:** Aylesse Sordillo, Cornelia I. Bargmann

## Abstract

Coordinated transitions between mutually exclusive motor states are central to behavioral decisions. During locomotion, the nematode *Caenorhabditis elegans* spontaneously cycles between forward runs, reversals, and turns with complex but predictable dynamics. Here we provide insight into these dynamics by demonstrating how RIM interneurons, which are active during reversals, act in two modes to stabilize both forward runs and reversals. By systematically quantifying the roles of RIM outputs during spontaneous behavior, we show that RIM lengthens reversals when depolarized through glutamate and tyramine neurotransmitters and lengthens forward runs when hyperpolarized through its gap junctions. RIM is not merely silent upon hyperpolarization: RIM gap junctions actively reinforce a hyperpolarized state of the reversal circuit. Additionally, the combined outputs of chemical synapses and gap junctions from RIM regulate forward-to-reversal transitions. Our results indicate that multiple classes of RIM synapses create behavioral inertia during spontaneous locomotion.

## INTRODUCTION

Neurons coordinate their activity across networks using a variety of signals: fast chemical transmitters, biogenic amines, neuropeptides, and electrical coupling via gap junctions (Tritsch & Sabatini, 2012; Zell et al., 2020; Taylor et al., 2019; P. Liu et al., 2017; Nagy et al., 2019). Signals from many neurons coalesce to generate large-scale brain activity patterns that are correlated with movement, while reflecting the animal’s memory, internal state, and sensory experience (Kato et al., 2015; Musall et al., 2019). The mechanisms for generating stable, mutually-exclusive activity and behavioral states across networks, while allowing behavioral flexibility, are incompletely understood.

The relationships between neurons, synapses, circuits, and behavior can be addressed precisely in the compact nervous system of *C. elegans.* Like many animals, *C. elegans* has locomotion-coupled, global brain activity states (Kato et al., 2015; Musall et al., 2019; Nguyen et al., 2016; Venkatachalam et al., 2016). Many of its integrating interneurons and motor neurons are active during one or more of three basic motor behaviors – forward runs, reversals, and turns (Figure 1A). A set of interneurons including AIB, AVA, and RIM are active whenever animals reverse (Gordus et al., 2015; Kato et al., 2015; Nguyen et al., 2016; Venkatachalam et al., 2016); a different set, AIY, RIB, and AVB, are active during forward runs (Kaplan et al., 2020; Kato et al., 2015; Li et al., 2014; Nguyen et al., 2016); and a set including AIB, RIB, and RIV are active during sharp omega turns, which typically follow a reversal (Kato et al., 2015; Nguyen et al., 2016; Venkatachalam et al., 2016; Wang et al., 2020). The functional role of each integrating neuron can be evaluated by considering the neuron’s regulation of specific locomotor features, like reversal speed or turn angle, and its influence on locomotor transitions. The AVA neurons, for example, are backward command neurons that drive reversals; when AVA neurons are optogenetically depolarized animals reverse, and when AVA neurons are ablated or acutely silenced reversals are short and infrequent (Chalfie et al., 1985; Gordus et al., 2015; P. Liu et al., 2017; Pokala et al., 2014; Roberts et al., 2016). Acute silencing of AVA often causes aberrant pauses – thwarted reversals – followed by a turn, indicating that AVA neurons are required for the execution of a reversal, but not for the global dynamics of the forward-reversal-turn sequence (Kato et al., 2015; Pokala et al., 2014). Other neurons in the locomotor circuit are implicated in transition dynamics. For example, altering AIB and RIB activity can change the probability and timing of the reversal-to-turn transition without generating abnormal pause states (Pokala et al., 2014, Wang et al., 2020).

**Figure 1.**
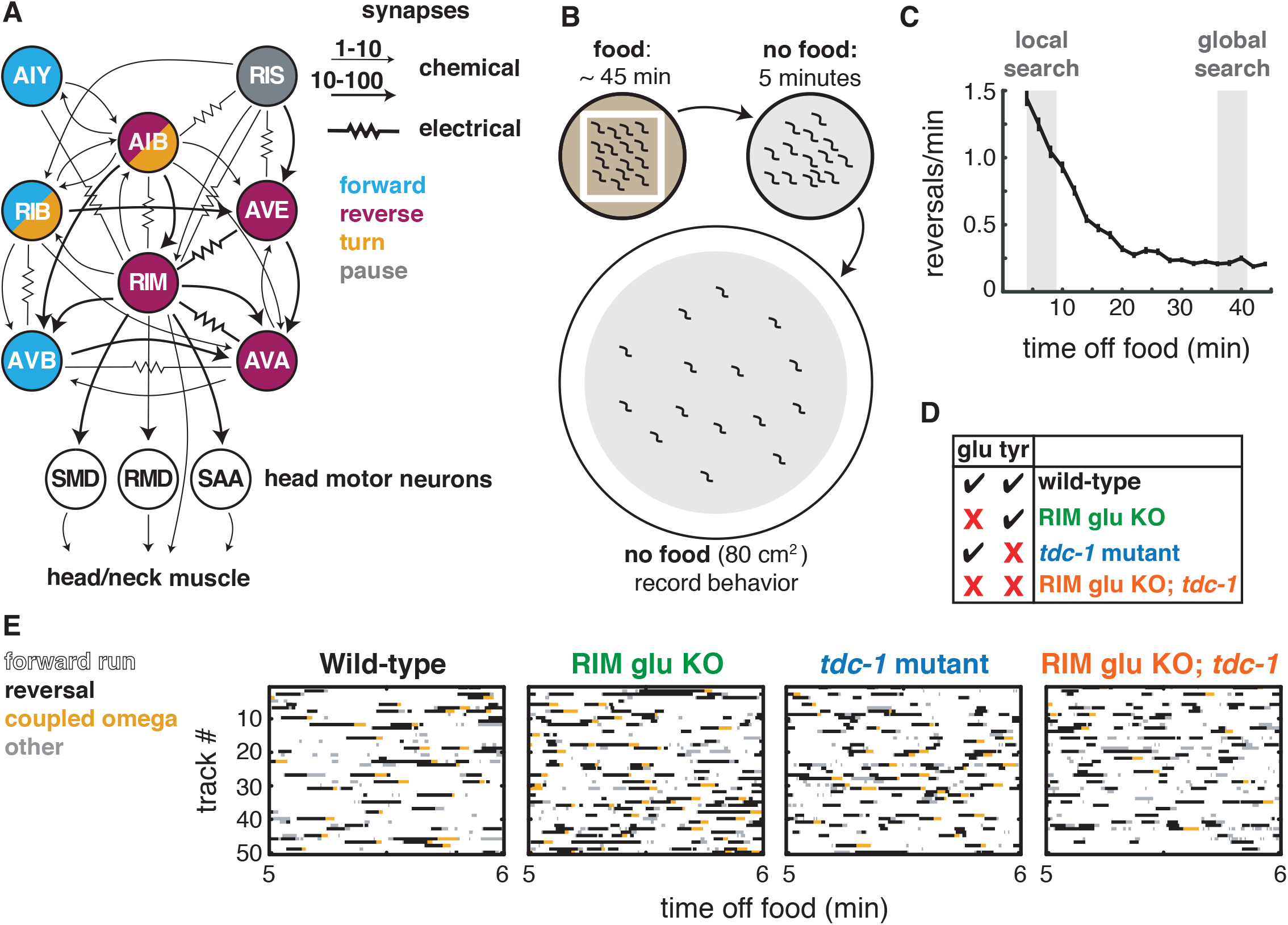
Two RIM neurotransmitters affect spontaneous locomotion. **(A)** RIM synapses with interneurons, motor neurons, and muscle implicated in spontaneous foraging behavior (Cook et al., 2019; White et al., 1986). Colors of neurons indicate associated locomotor states based on neural manipulations and functional calcium imaging (Alkema et al., 2005; Gray et al., 2005; Kato et al., 2015; Li et al., 2014; Pokala et al., 2014; Steuer Costa et al., 2019; Wang et al., 2020; Zheng et al., 1999). **(B)** Off-food foraging assay. **(C)** Mean reversals per minute of wild-type animals in foraging assays. Vertical lines indicate standard error of the mean. Gray shaded boxes indicate local search (4-9 min off food) and global search (36-41 min off food) intervals analyzed in subsequent figures. n = 324. **(D)** RIM neurotransmitter mutants. RIM glu KO = RIM-specific knockout of the vesicular glutamate transporter EAT-4 (Figure 1–figure supplement 1). *tdc-1,* tyrosine decarboxylase mutant, which lacks tyramine in RIM and octopamine in RIC. **(E)** Ethograms of 50 randomly chosen tracks per genotype during minute 5-6 of local search. Color code: white, forward runs; black, reversals; yellow, omega turns coupled to a reversal; grey, pauses, shallow turns, and omega turns that were not preceded by a reversal.

Among the interneurons in the locomotor circuit, RIM, a pair of motor/interneurons, has both straightforward and apparently paradoxical functions (Figure 1A). RIM is active during both spontaneous and stimulus-evoked reversals, and its activity correlates with reversal speed (Gordus et al., 2015; Kagawa-Nagamura et al., 2018; Kato et al., 2015). RIM releases the neurotransmitter tyramine, which extends reversals by inhibiting the AVB forward-active neurons and suppresses head oscillations by inhibiting the head muscles, in both cases through the tyramine-gated chloride channel LGC-55 (Alkema et al., 2005; Pirri et al., 2009). RIM tyramine also sharpens reversal-coupled omega turns by activating SER-2, a G protein-coupled receptor on motor neurons (Donnelly et al., 2013). In addition to these effects on locomotion parameters, RIM has puzzling effects on behavioral transitions. Optogenetic depolarization of RIM drives reversals, but ablation of RIM paradoxically increases spontaneous reversals, indicating that RIM can either induce or suppress reversals (Gordus et al., 2015; Gray et al., 2005; Guo et al., 2009; Lopez-Cruz et al., 2019; Zheng et al., 1999). RIM also mediates competition between sensory inputs and motor circuits, generating variability in behavioral responses to external stimuli (Gordus et al., 2015; Ji et al., 2019), and biases choices between attractive and aversive stimuli (Ghosh et al., 2016; Hapiak et al., 2013; Li et al., 2012; Wragg et al., 2007). On longer timescales, RIM modulates learning as well as physiological responses to temperature or unfolded protein stress (De Rosa et al., 2019; Fu et al., 2018; Ha et al., 2010; Jin et al., 2016; Ozbey et al., 2020).

Here we develop an integrated view of RIM’s role in locomotor features, motor transitions, and behavioral dynamics through cell-specific manipulation of its synapses. In addition to tyramine, RIM expresses the vesicular glutamate transporter EAT-4, identifying it as one of the 38 classes of glutamatergic neurons in *C. elegans* (Lee et al., 1999; Serrano-Saiz et al., 2013). RIM also forms gap junctions with multiple neurons whose activity is associated with reversals (AIB, AVA, AVE), as well as neurons active during pauses (RIS) and forward runs (AIY) (Cook et al., 2019; White et al., 1986)(Figure 1A). *eat-4* and gap junction subunits are broadly expressed throughout the foraging circuit, precluding a simple interpretation of null mutants in these genes (Bhattacharya et al., 2019; Serrano-Saiz et al., 2013). Therefore, we used a cell-specific knockout of *eat-4* and a cell-specific gap junction knockdown to isolate the synaptic functions of RIM. By examining behavioral effects of multiple transmitters and gap junctions, we reveal distinct functions of RIM during reversals, when its activity is high, and during forward locomotion, when its activity is low. Notably, our results indicate that while RIM depolarization extends reversals, the propagation of hyperpolarization through RIM gap junctions extends the opposing forward motor state. This work indicates that a single interneuron class employs different classes of synapses to shape mutually exclusive behaviors.

## RESULTS

### RIM glutamate and tyramine suppress spontaneous reversals and increase reversal length

The goal of this work was to understand how RIM influences spontaneous behavioral dynamics, including individual features of locomotion and transitions between motor states. We used an off-food foraging assay in which forward, reversal, and turn behaviors emerge from predictable internal states (Calhoun et al., 2014; Gray et al., 2005; Hills et al., 2004; Lopez-Cruz et al., 2019; Wakabayashi et al., 2004) (Figure 1B). When removed from food and placed on a featureless agar surface, *C. elegans* engages in local search, in which a high frequency of spontaneous reversals limits dispersal from the recently encountered food source. Over about fifteen minutes, animals spontaneously transition into global search, a state with infrequent reversals and long forward runs that promotes dispersal (Figure 1C). We recorded animals throughout this assay, and identified and quantified reversals, turns, forward runs, and pauses from behavioral sequences (example tracks in Figure 1E). The full dataset is available for further analysis (https://github.com/BargmannLab/SordilloBargmann2021), and a summary of results is included in Figure 8.

We began by examining the effects of RIM glutamate on local search (Figure 1D-E). *C. elegans* mutants lacking the vesicular glutamate transporter *eat-4* or various glutamate receptors have abnormal local search behaviors (Baidya et al., 2014; Chalasani et al., 2007; Choi et al., 2015; Hills et al., 2004; Lopez-Cruz et al., 2019). To selectively inactivate glutamatergic transmission from RIM, we used an FRT-flanked endogenous *eat-4* locus and expressed FLP recombinase under a *tdc-1* promoter, which intersects with *eat-4* only in RIM (Lopez-Cruz et al., 2019) (Figure 1–figure supplement 1). The resulting animals lacking RIM glutamate had an increased frequency of reversals during local search, but not global search (Figure 1E, 2A-B; Figure 2– figure supplement 1).

**Figure 2.**
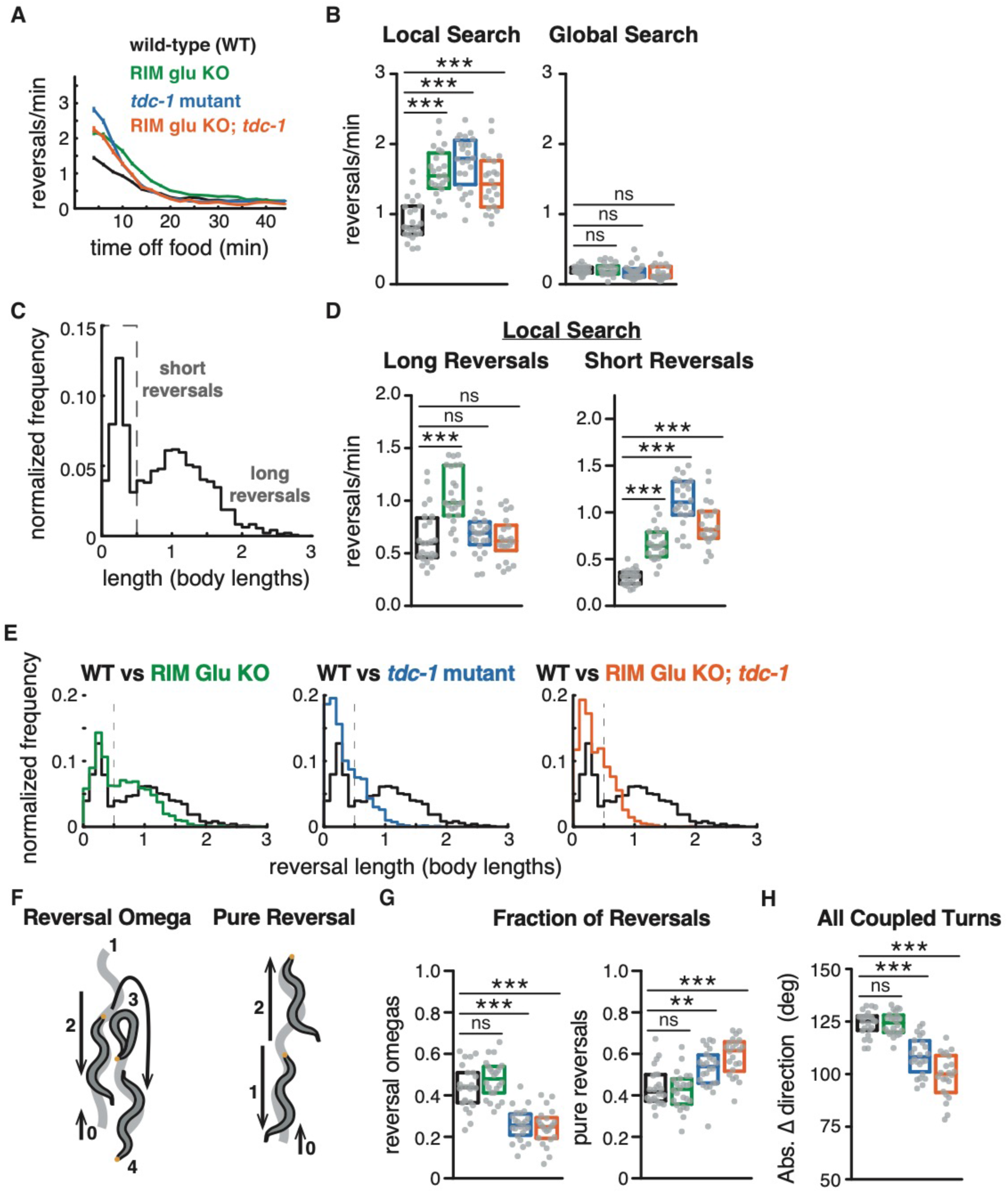
RIM glutamate and tyramine suppress spontaneous reversals and increase reversal length. **(A)** Mean reversals per minute in foraging assays for genotypes analyzed in Figures 2-3. Vertical dashes indicate standard error of the mean. n = 296-332. All strains bear *tdc-1p*::nFLP and the *elt-2p*::nGFP marker (Figure 2–figure supplement 1, Supplementary file 1). **(B)** Mean frequency of all reversals during local search (4-9 min off food, left) and global search (36-41 min off food, right). **(C)** Normalized probability distribution of wild-type reversal lengths during local search. Short reversals cover less than 0.5 body lengths. **(D)** Mean frequency of long reversals (>0.5 body lengths, left) and short reversals (<0.5 body lengths, right) during local search. **(E)** Normalized probability distribution of mutant reversal lengths during local search, plotted with WT distributions. **(F)** A forward moving animal (0) initiates a reversal (1-2) that is coupled to an omega turn (3) and terminates in forward movement (4) (reversal omega, left). A forward moving animal (0) initiates a reversal (1) that terminates in forward movement (2) (pure reversal, right). Yellow dot indicates nose. **(G)** Fraction of all reversals during local search that terminate in an omega turn (left) or forward movement (right) for each genotype. **(H)** Absolute change in direction after a reversal-turn maneuver (including omega and shallower turns) for each genotype. **(B, D, G, H)** Each gray dot is the mean for 12-15 animals on a single assay plate, with 22-24 plates per genotype. Boxes indicate median and interquartile range for all assays. Asterisks indicate statistical significance compared to WT using a Kruskal-Wallis test with Dunn’s multiple comparisons test (** = p-value < 0.01, *** = p-value < 0.001, ns = p-value ≥ 0.05). In **(C, E)** n=1443-2760 events per genotype. The reversal defects in (RIM) tyramine- and (RIC) octopamine-deficient *tdc-1* mutants are not shared by octopamine-deficient *tbh-1* mutants (Figure 2–figure supplement 2).

RIM is the primary neuronal source of tyramine, whose synthesis requires the tyrosine decarboxylase encoded by *tdc-1* (Alkema et al., 2005). As previously reported, *tdc-1* mutants had an increased reversal frequency during local search (Figure 1E, 2A-B)(Alkema et al., 2005). *tdc-1* is also expressed in RIC neurons, where it is used, together with *tbh-1,* in the biosynthesis of the neurotransmitter octopamine (Alkema et al., 2005). *tbh-1* did not affect reversal frequency during local search, identifying tyramine as the relevant transmitter for reversals (Figure 2–figure supplement 2). The RIM glu KO; *tdc-1* double mutant was similar to each single mutant (Figure 2A-B). Thus, both of RIM’s neurotransmitters, glutamate and tyramine, suppress spontaneous reversals.

Reversals during local search segregate into short reversals of less than half a body length, and long reversals that average >1 body length (Gray et al., 2005) (Figure 2C-D). Using these criteria, both short and long reversals increased in frequency in RIM glu KO animals during local search, but only short reversals increased in frequency in *tdc-1* mutants or the RIM glu KO; *tdc-1* double mutant (Figure 2D). To better understand this distinction, we conducted an analysis of the full reversal length distribution (Figure 2E). In fact, both RIM glu KO animals and *tdc-1* mutants had decreased reversal lengths compared to wild-type, with a stronger effect in *tdc-1* mutants, indicating that RIM glutamate and tyramine both extend reversal length.

Long reversals are more likely to be followed by an omega turn than short reversals (Chalasani et al., 2007; Croll, 1975; Gray et al., 2005; Huang et al., 2006; Wang et al., 2020; Zhao et al., 2003) (Figure 2F). The fraction of reversal omegas was reduced in *tdc-1* mutants (Figure 2G, left), while pure reversals that terminate in an immediate forward run increased (Figure 2G, right). As previously reported, omega turn angles were shallower in *tdc-1* mutants (Figure 2H). By contrast, RIM glu KO animals had normal reversal-omega frequencies and turn angles after reversals, despite a decrease in reversal length (Figure 2G-H).

### RIM neurotransmitters distinguish reversal and reversal-omega behaviors

Analysis of the frequency distributions of all reversal lengths, speeds, and durations uncovered additional distinctions between the functions of RIM glutamate and tyramine (Figure 3A-I). First, while reversal lengths were decreased in a graded fashion by RIM glu KO or *tdc-1* (Figure 2E, 3A), reversal speeds were substantially reduced only in *tdc-1* mutants (Figure 3B). *tdc-1* and RIM glu KO had similar decreases in reversal durations (Figure 3C). We found that genetic markers and background controls could affect these parameters by up to 12%; with that in mind, we interpret only effect sizes of ≥0.15 in these and other quantitative experiments (see Materials and methods and Figure 2–figure supplement 1).

**Figure 3.**
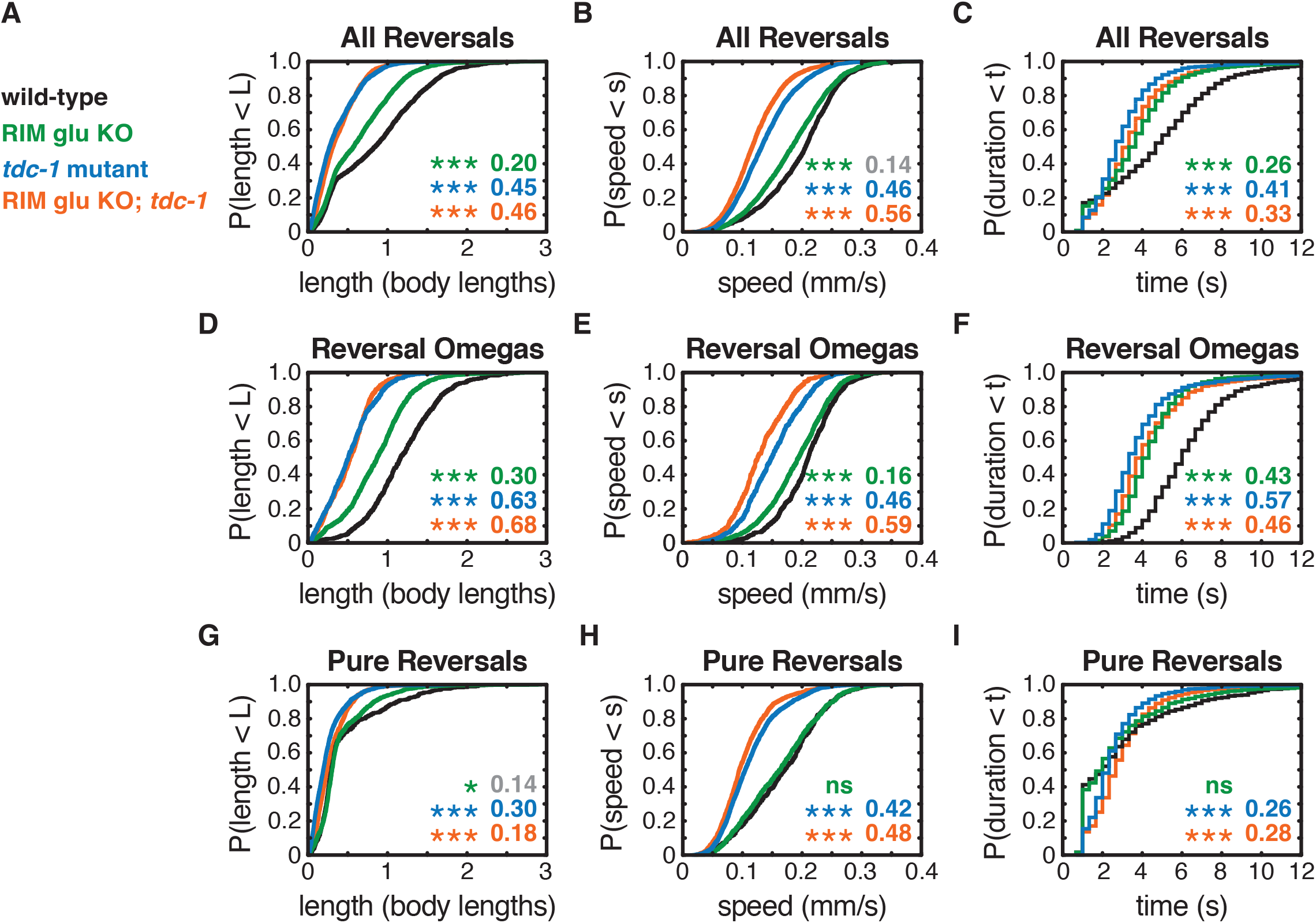
RIM neurotransmitters differently affect reversal and reversal-omega behaviors. **(A-C)** For all reversals during local search, empirical cumulative distributions of reversal length **(A)** reversal speed **(B)** and reversal duration **(C)**. **(D-F)** For reversal-omega maneuvers during local search, empirical cumulative distributions of reversal length **(D)** reversal speed **(E)** and reversal duration **(F)**. **(G-I)** For pure reversals during local search, empirical cumulative distributions of reversal length **(G)** reversal speed **(H)** and reversal duration **(I)**. Asterisks indicate statistical significance compared to WT using a two-sample Kolmogorov Smirnov test, with a Bonferroni correction (* = p-value < 0.05, *** = p-value < 0.0001, ns = p-value ≥ 0.05). Numbers in figures indicate effect size (See Materials and methods). Although statistically significant, the smaller effect sizes indicated in gray are similar to values from control strains (e.g., Figure 2–figure supplement 1), and fall under the 0.15 cutoff for interpretation established from those controls. For each value, n = 500-3132 events from 22-24 assays, 12-15 animals per assay (Supplementary file 3). The reversal defects in (RIM) tyramine- and (RIC) octopamine-deficient *tdc-1* mutants are not shared by octopamine-deficient *tbh-1* mutants (Figure 2–figure supplement 2).

Separating different classes of reversals (Figure 2F-G) revealed that the RIM glu KO decreased reversal-omega duration but did not affect pure reversal duration (Figure 3D-I). *tdc-1* mutants decreased the duration of reversal omegas, increased the duration of pure reversals, and decreased the speed of all reversals (Figure 3D-I). RIM glu KO; *tdc-1* double mutant animals resembled *tdc-1* mutants.

Forward runs are heterogeneous compared to reversals, with an exponential distribution of durations (Figure 2–figure supplement 2) (Wakabayashi et al., 2004). Neither RIM glu KO animals nor *tdc-1* mutants appreciably decreased forward run durations compared to controls (Figure 2–figure supplement 1, Figure 2–figure supplement 2). Both *tdc-1* and *tbh-1* mutants had substantially diminished forward speeds, suggesting a role of octopamine in forward locomotion (Figure 2–figure supplement 2). Because the octopaminergic RIC neurons were not the focus of this work, forward speed was not examined further.

In summary, tyramine affects both the speed and the duration of all classes of reversals, whereas RIM glutamate only increases the duration of reversals that are coupled to omega turns. RIM neurotransmitters do not substantially affect forward run durations, consistent with low RIM activity during forward runs. RIC octopamine increases forward speed.

### Additional RIM transmitters contribute to global search dynamics

In addition to glutamate and tyramine, RIM expresses multiple neuropeptides (*flp-18, pdf-2*, and others) (Bhardwaj et al., 2018; Ghosh et al., 2016; Taylor et al., 2019). Release of both classical transmitters and neuropeptides is inhibited by the tetanus toxin light chain, which cleaves the synaptic vesicle fusion protein synaptobrevin (Schiavo et al., 1992). Expression of tetanus toxin in RIM and RIC under the *tdc-1* promoter resulted in defects resembling those of *tdc-1* mutants (Figure 4A-C): reversal frequency increased, while reversal length, speed, and durations decreased, during local search behavior (Figure 4D-G). Efficient RIM-only promoters are not available, but expression of tetanus toxin in RIC alone yielded minor defects in reversal frequency and speed, implicating RIM in the regulation of reversal parameters (Figure 4–figure supplement 1, Figure 4–figure supplement 2). RIC tetanus toxin expression reduced forward locomotion speed to a similar extent as *tbh-1* mutants (Figure 4–figure supplement 1).

**Figure 4.**
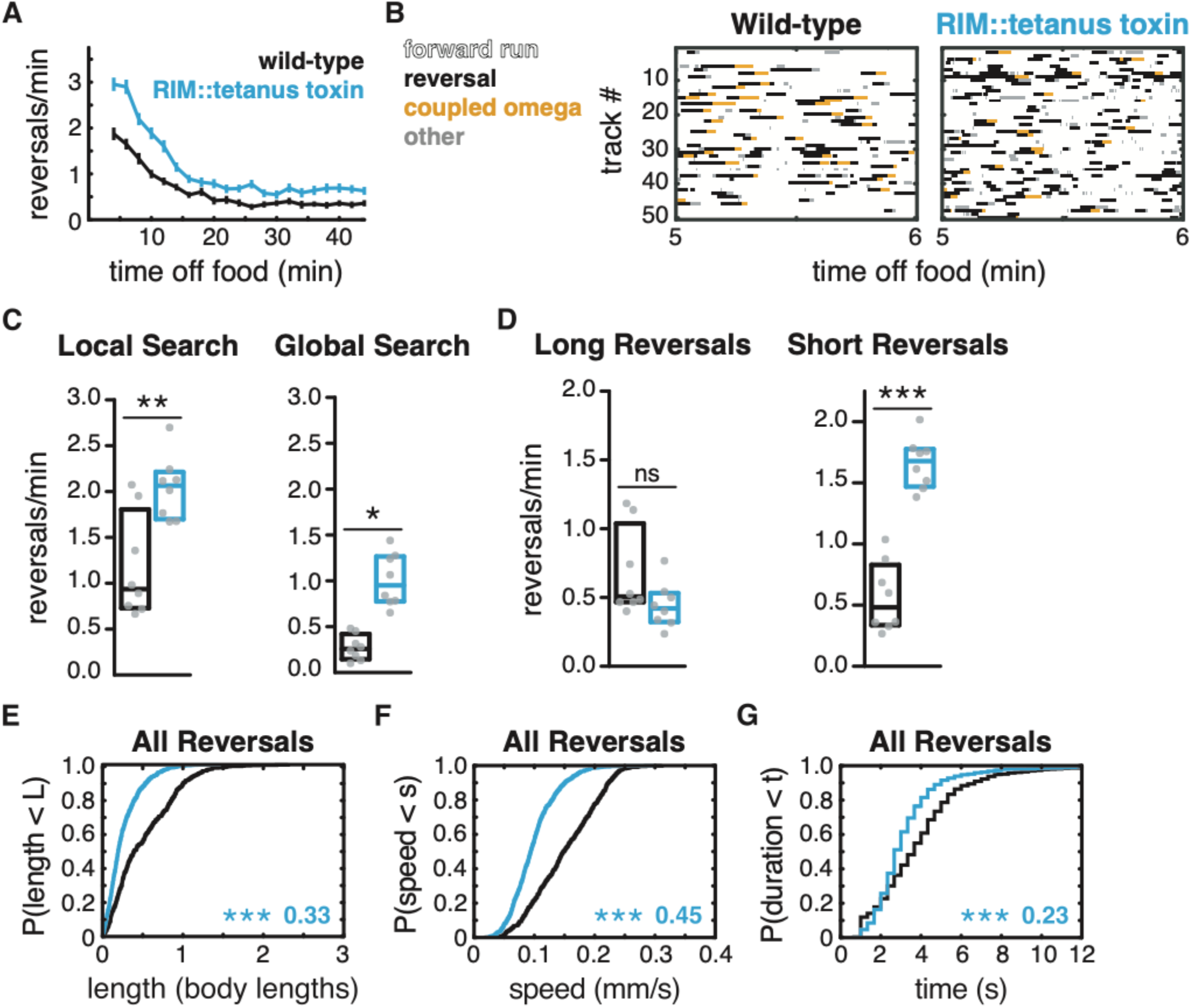
Additional RIM transmitters contribute to global search dynamics. **(A)** Mean reversals per minute in animals expressing tetanus toxin light chain under the RIM-and RIC-specific *tdc-1* promoter. Defects are milder or absent when tetanus toxin is expressed under the RIC-specific *tbh-1* promoter (Figure 4–figure supplement 1). Vertical dashes indicate standard error of the mean. n = 103-111. **(B)** Ethograms of 50 randomly chosen tracks per genotype during minute 5-6 of local search. Color code: white, forward runs; black, reversals; yellow, omega turns coupled to a reversal; grey, pauses, shallow turns, and omega turns that were not preceded by a reversal. **(C)** Mean frequency of all reversals during local search (4-9 min off food, left) and global search (36-41 min off food, right). **(D)** Mean frequency of long reversals (>0.5 body lengths, left) and short reversals (<0.5 body lengths, right) during local search. **(E-G)** For all reversals during local search, empirical cumulative distributions of reversal length **(E)** reversal speed **(F)** and reversal duration **(G)**. **(C-D)** Each gray dot is the mean for 12-15 animals on a single assay plate. Boxes indicate median and interquartile range for all assays. Asterisks indicate statistical significance compared to WT using a Mann-Whitney test (*=p-value < 0.05, ** = p-value < 0.01, *** = p-value < 0.001, ns = p-value ≥ 0.05). **(E–G**) Asterisks indicate statistical significance compared to WT using a two-sample Kolmogorov Smirnov test (*** = p-value < 0.0001). Numbers indicate effect size. For each value, n = 595-1066 events from 8 assays, 12-15 animals per assay (Supplementary file 3).

The expression of tetanus toxin in RIM and RIC also increased reversals during the global search period, an effect that was not observed in RIM glutamate KO or tyramine-deficient mutants (Figure 4A,C). Tetanus toxin expression in RIC alone did not affect global search (Figure 4–figure supplement 1). These results suggest that a synaptobrevin-dependent transmitter from RIM, perhaps a neuropeptide, suppresses reversals during global search.

### Artificial hyperpolarization of RIM reveals unexpected functions in forward runs

To relate RIM functions to its membrane potential, we hyperpolarized RIM by expressing the *Drosophila* histamine-gated chloride channel (HisCl) under the *tdc-1* promoter and exposing the animals to histamine while off food (Pokala et al., 2014) (Figure 5A-B). Unexpectedly, silencing RIM with HisCl led to a substantial decrease in spontaneous reversal frequency and a striking increase in forward run duration, both in wild-type and in *tdc-1* mutants (Figure 5C-D). The effects on reversal frequency were opposite to those of RIM ablation, RIM neurotransmitter mutants, or RIM::tetanus toxin expression, all of which increased spontaneous reversal frequency (Alkema et al., 2005; Gray et al., 2005) (Figures 2-4). The opposite effects of RIM ablation and acute silencing suggests that RIM has active functions when hyperpolarized that stabilize and extend forward runs.

**Figure 5.**
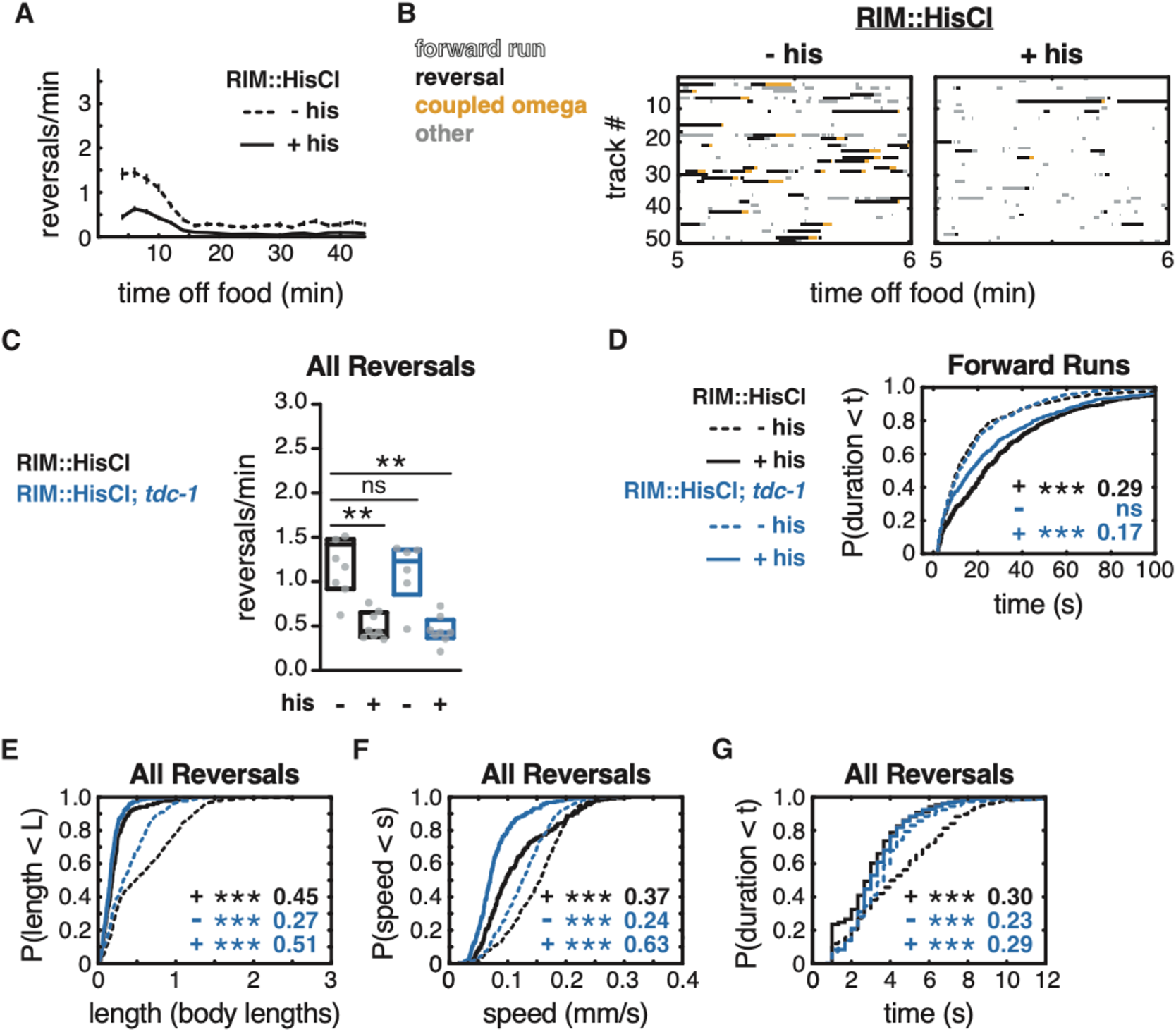
Artificial hyperpolarization of RIM extends forward runs and suppresses reversals. **(A)** Mean reversals per minute in animals expressing HisCl in RIM, with (+ his) or without (- his) histamine treatment. Vertical dashes indicate standard error of the mean. n = 93-109. **(B)** Ethograms of 50 randomly chosen tracks per genotype during minute 5-6 of local search. Color code: white, forward runs; black, reversals; yellow, omega turns coupled to a reversal; grey, pauses, shallow turns, and omega turns that were not preceded by a reversal. **(C)** Mean frequency of all reversals during local search, with or without histamine, in wild-type or *tdc-1* animals expressing HisCl in RIM. Each gray dot is the mean for 12-15 animals on a single assay plate. Boxes indicate median and interquartile range for all assays. Asterisks indicate statistical significance compared to WT untreated controls using a Kruskal-Wallis test with Dunn’s multiple comparisons test (**=p-value < 0.01, ns = p-value ≥ 0.05). **(D)** Durations of forward runs during local search with (solid lines) and without (dashed lines) histamine, in wild-type or *tdc-1* animals expressing HisCl in RIM; empirical cumulative distributions include all runs ≥ 2 s. **(E-G)** For all reversals during local search, empirical cumulative distributions of reversal length **(E)** reversal speed **(F)** and reversal duration **(G)** with or without histamine, in wild-type or *tdc-1* animals expressing HisCl in RIM. **(C, E-G)** Asterisks indicate statistical significance compared to WT untreated controls using a two-sample Kolmogorov Smirnov test with a Bonferroni correction (*** = p-value < 0.0001, ns = p-value ≥ 0.05). Numbers indicate effect size. n = 251–579 reversals from 6–8 assays, 12–15 animals per assay (Supplementary file 3).

Reversal length, speed, and duration were greatly reduced by hyperpolarization of RIM, effects that were similar to but stronger than the effect of *tdc-1* or tetanus toxin (Figure 5E-G). Introducing RIM::HisCl into a *tdc-1* mutant yielded the stronger reversal defects of the RIM::HisCl strain (Figure 5E-G). These results suggest that RIM glutamate and tyramine are released when RIM is depolarized, as expected, to extend reversals and increase reversal speed. However, the stronger effects of RIM::HisCl indicate that hyperpolarization affects other targets as well.

### RIM gap junctions stabilize forward runs

To explain the effect of hyperpolarized RIM neurons, we considered the gap junctions that RIM forms with a variety of other neurons in the local search circuit (Figure 1A). RIM shares the most gap junctions with AVA and AVE which, like RIM, have high activity during reversals and low activity during forward runs. We hypothesized that RIM gap junctions stabilize the forward motor state by propagating hyperpolarizing currents between RIM and AVA (and possibly other) neurons, thereby preventing their depolarization.

Invertebrate gap junctions are made up of innexin subunits, which assemble as homo- or heteromers of eight subunits on each of the two connected cells (Burendei et al., 2020; Oshima et al., 2016). Most *C. elegans* neurons express multiple innexin genes; RIM neurons express eleven innexin genes, including *unc-9* (Bhattacharya et al., 2019). *unc-9* is expressed in many classes of neurons, and mutants have strong defects in forward and backward locomotion (Brenner, 1974; Kawano et al., 2011; P. Liu et al., 2017; Q. Liu et al., 2006; Park & Horvitz, 1986; Shui et al., 2020; Starich et al., 2009). To bypass its broad effects, neuronal *unc-9* function can be reduced in a cell-specific fashion by expressing UNC-1(n494), a dominant negative allele of a stomatin-like protein that is an essential component of neuronal UNC-9 gap junctions (Chen et al., 2007; Jang et al., 2017). We knocked down UNC-9 gap junctions in RIM by driving *unc-1(n494)* cDNA under the *tdc-1* promoter. While unlikely to inactivate all RIM innexins and gap junctions, this manipulation should alter *unc-9* gap junction signaling in a RIM-selective manner.

RIM gap junction knockdown animals had superficially coordinated locomotion and exhibited the characteristic shift from local to global search over time (Figure 6A). However, these gap junction knockdown animals had a greatly increased frequency of reversals compared to wild-type (Figure 6B). Both short and long reversals were increased in frequency during both local search and global search, while reversal length, speed, and duration were largely unaffected (Figure 6C-G, Figure 6–figure supplement 1). The RIM gap junction knockdown also resulted in a substantial decrease in forward run duration (Figure 6H, Figure 6–figure supplement 1). Combining the gap junction knockdown with *tdc-1* mutation yielded additive effects, with both forward and reversal parameters altered (Figure 6–figure supplement 2). These results support the hypothesis that *unc-9*-containing gap junctions in RIM promote forward locomotion.

**Figure 6.**
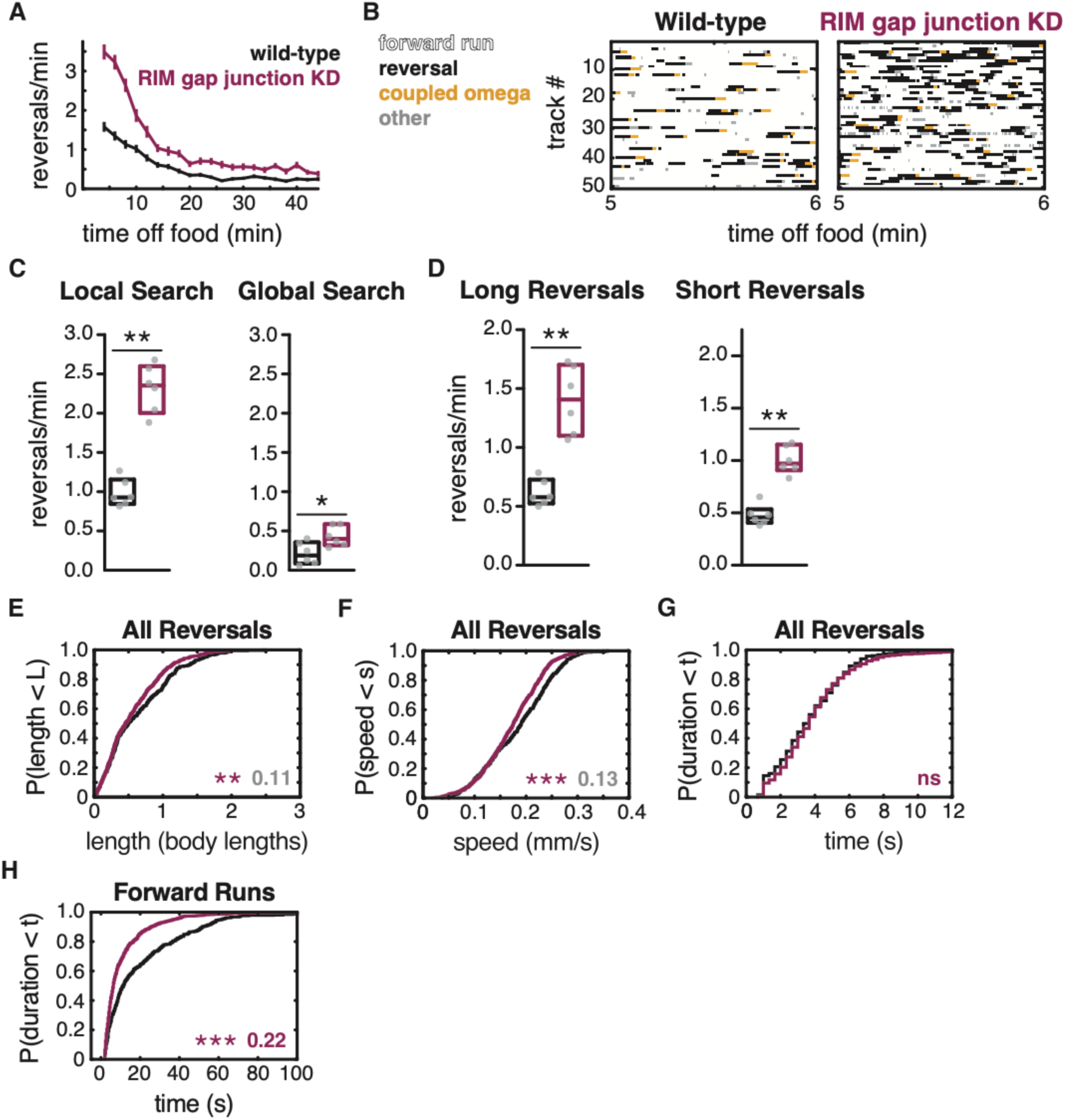
RIM gap junctions stabilize forward runs. **(A)** Mean reversals per minute in animals bearing an *unc-1(n494)* dominant negative transgene to knock down *unc-9-*containing gap junctions (RIM gap junction KD; note that the *tdc-1* promoter also expresses *unc-1(n494)* in RIC). Vertical dashes indicate standard error of the mean. n = 77-85. **(B)** Ethograms of 50 randomly chosen tracks per genotype during minute 5-6 of local search. Color code: white, forward runs; black, reversals; yellow, omega turns coupled to a reversal; grey, pauses, shallow turns, and omega turns that were not preceded by a reversal. **(C)** Mean frequency of all reversals during local search (4-9 min off food, left) and global search (36-41 min off food, right). **(D)** Mean frequency of long reversals (>0.5 body lengths, left) and short reversals (<0.5 body lengths, right) during local search. **(E-G)** For all reversals during local search, empirical cumulative distributions of reversal length **(E)** reversal speed **(F)** and reversal duration **(G)**. **(H)** Durations of forward runs during local search; empirical cumulative distributions include all runs ≥ 2 s. **(C-D)** Each gray dot is the mean for 12-15 animals on a single assay plate. Boxes indicate median and interquartile range for all assays. Asterisks indicate statistical significance compared to WT using a Mann-Whitney test (* =p-value < 0.05, ** =p-value < 0.01). **(E-H**) Asterisks indicate statistical significance compared to WT using a two-sample Kolmogorov Smirnov test (**=p-value < 0.01, *** = p-value < 0.001, ns = p-value ≥ 0.05). Numbers indicate effect size. Although statistically significant, the smaller effect sizes indicated in gray are similar to values from control strains (e.g., Figure 2–figure supplement 1). n = 330-933 reversals from 6 assays, 12-15 animals per assay (Supplementary file 3).

To ask whether the *unc-9-*containing gap junctions propagate the effects of RIM hyperpolarization, we crossed the RIM gap junction knockdown into the RIM::HisCl strain. Combining the RIM gap junction knockdown with RIM hyperpolarization resulted in mutual suppression of their effects, so that double mutants had a similar reversal frequency and forward run duration to wild-type animals (Figure 7A-C). These results suggest that forward states are stabilized during RIM hyperpolarization in part through *unc-9-*containing gap junctions (Figure 7C, Figure 7–figure supplement 1).

**Figure 7.**
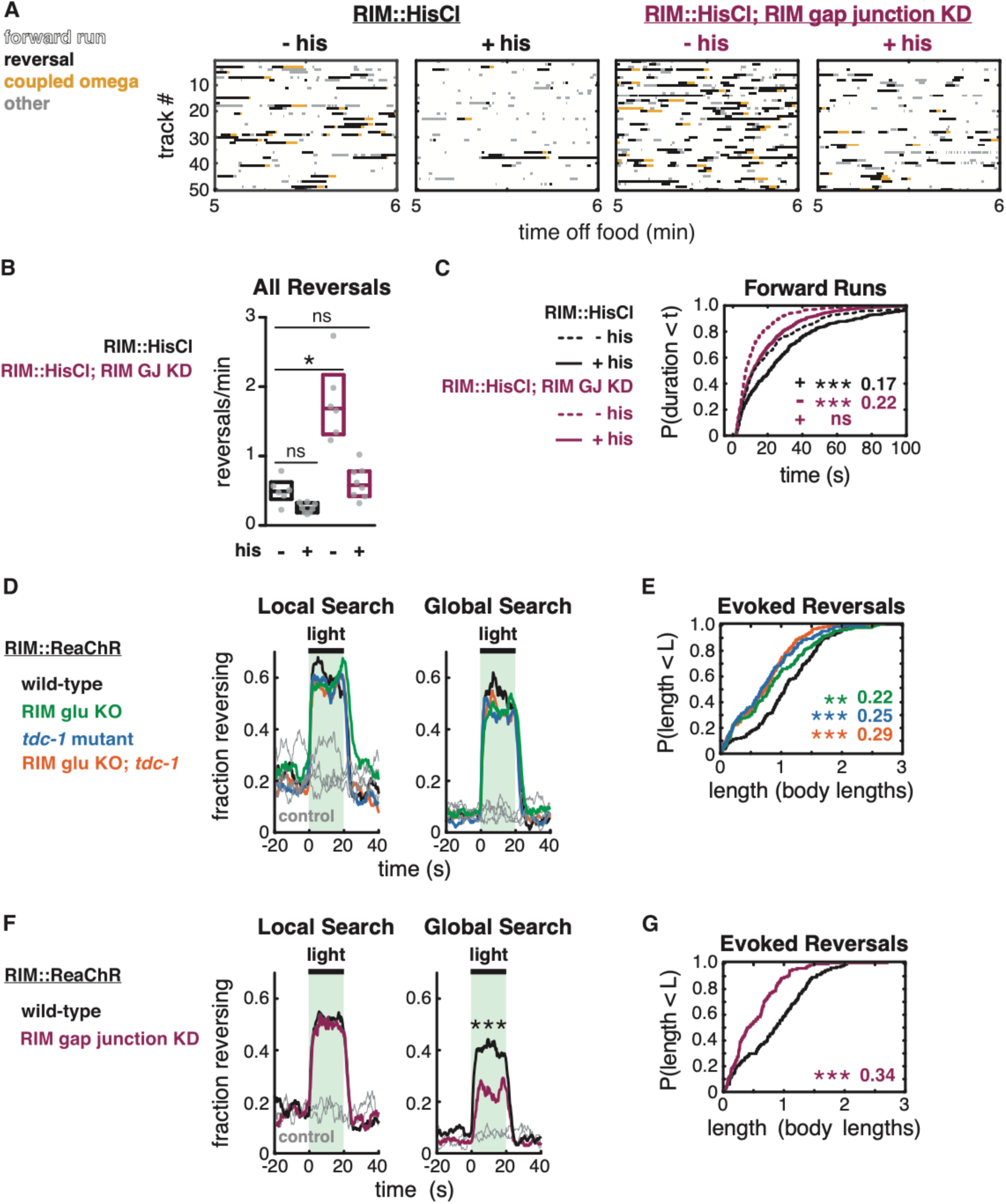
RIM gap junctions mediate effects of hyperpolarization and depolarization. **(A-C)** Behavior of RIM::HisCl and RIM::HisCl; RIM gap junction knockdown animals. **(A)** Ethograms of 50 randomly chosen tracks per genotype during minute 5-6 of local search. Color code: white, forward runs; black, reversals; yellow, omega turns coupled to a reversal; grey, pauses, shallow turns, and omega turns that were not preceded by a reversal. **(B)** Mean frequency of all reversals during local search (4-9 min off food), with or without histamine. Each gray dot is the mean for 12-15 animals on a single assay plate. Boxes indicate median and interquartile range for all assays. Asterisks indicate statistical significance compared to untreated WT controls using a Kruskal-Wallis test with Dunn’s multiple comparisons test (* = p-value < 0.05, ns = p-value ≥ 0.05). **(C)** Durations of forward runs during local search with (solid lines) and without (dashed lines) histamine; empirical cumulative distributions include all runs ≥ 2 s. n = 330-819 events from 6-8 assays, 12-15 animals per assay. Asterisks indicate statistical significance compared to WT untreated controls using a two-sample Kolmogorov Smirnov test, corrected for multiple comparisons (*** = p-value < 0.0001, ns = p-value ≥ 0.05). Numbers indicate effect size. **(D-G)** Effects of RIM::ReaChR activation in wild-type, RIM glu KO, *tdc-1* mutants, RIM glu KO; *tdc-1* double mutants, and RIM gap junction knockdown animals. **(D,F)** Animals were exposed to light for 20 s (green shading), with or without all-trans retinal pre-treatment, during local search (8-14 min off food, left) or global search (38-44 min off food, right). Neurotransmitter mutants do not suppress optogenetically-evoked reversals **(D)**. RIM gap junction knockdown suppresses optogenetically-evoked reversals during global search **(F)** (*** p< 0.001, Figure 7–figure supplement 2). Similar results were obtained at lower and higher light levels. **(E,G)** For all reversals induced during the light pulse during local search (8-14 min off food), empirical cumulative distributions of reversal length. All animals were pre-treated with all-trans retinal. n = 119-193 reversals from 14-15 assays, 12-15 animals per assay, 2 **(E)** or 3 **(G)** light pulses per assay conducted 8-14 minutes after removal from food (Supplementary file 3). Asterisks indicate statistical significance compared to controls of the same genotype using a two-sample Kolmogorov Smirnov test with a Bonferroni correction (** = p-value < 0.01, *** = p-value < 0.001). Numbers indicate effect size.

### Strong depolarization of RIM engages neurotransmitter-independent functions

Optogenetic depolarization of RIM rapidly increases reversal frequency (Gordus et al., 2015; Guo et al., 2009; Lopez-Cruz et al., 2019)(Figure 7D). The frequency of optogenetically-induced reversals was unaffected by *tdc-1,* RIM glu KO, or the double mutant, whether examined during local search or during global search (Figure 7D, Figure 7–figure supplement 2). This result suggests that RIM does not require tyramine or glutamate neurotransmitters to trigger optogenetically-induced reversals.

We considered whether RIM gap junctions might propagate optogenetic depolarization to AVA command neurons. The RIM gap junction knockdown did not affect optogenetically-induced reversal frequencies during local search, but it did decrease optogenetically-induced reversals during global search (Figure 7F, Figure 7–figure supplement 2). These results suggest a minor role for *unc-9* gap junctions in the initiation of optogenetically-induced reversals.

Optogenetically-induced reversals were shorter in RIM glu KO, tyramine-deficient, and RIM gap junction knockdown animals than in wild-type (Figure 7E, G). Thus, optogenetically-induced reversals are extended by all RIM synaptic outputs.

## DISCUSSION

A cycle of forward runs interrupted by reversals and turns dominates the spontaneous locomotion of *C. elegans* during local search. We show here that RIM generates behavioral inertia to inform the dynamics of these locomotor states (Figure 8). RIM stabilizes reversals through its chemical synapses while depolarized and stabilizes forward runs through its gap junctions while hyperpolarized. Together with other results (Kawano et al., 2011), our results suggest that hyperpolarization through gap junctions is a recurrent circuit motif in *C. elegans* locomotion. In addition, our results indicate that RIM synapses jointly inhibit the forward-to-reversal transition.

**Figure 8.**
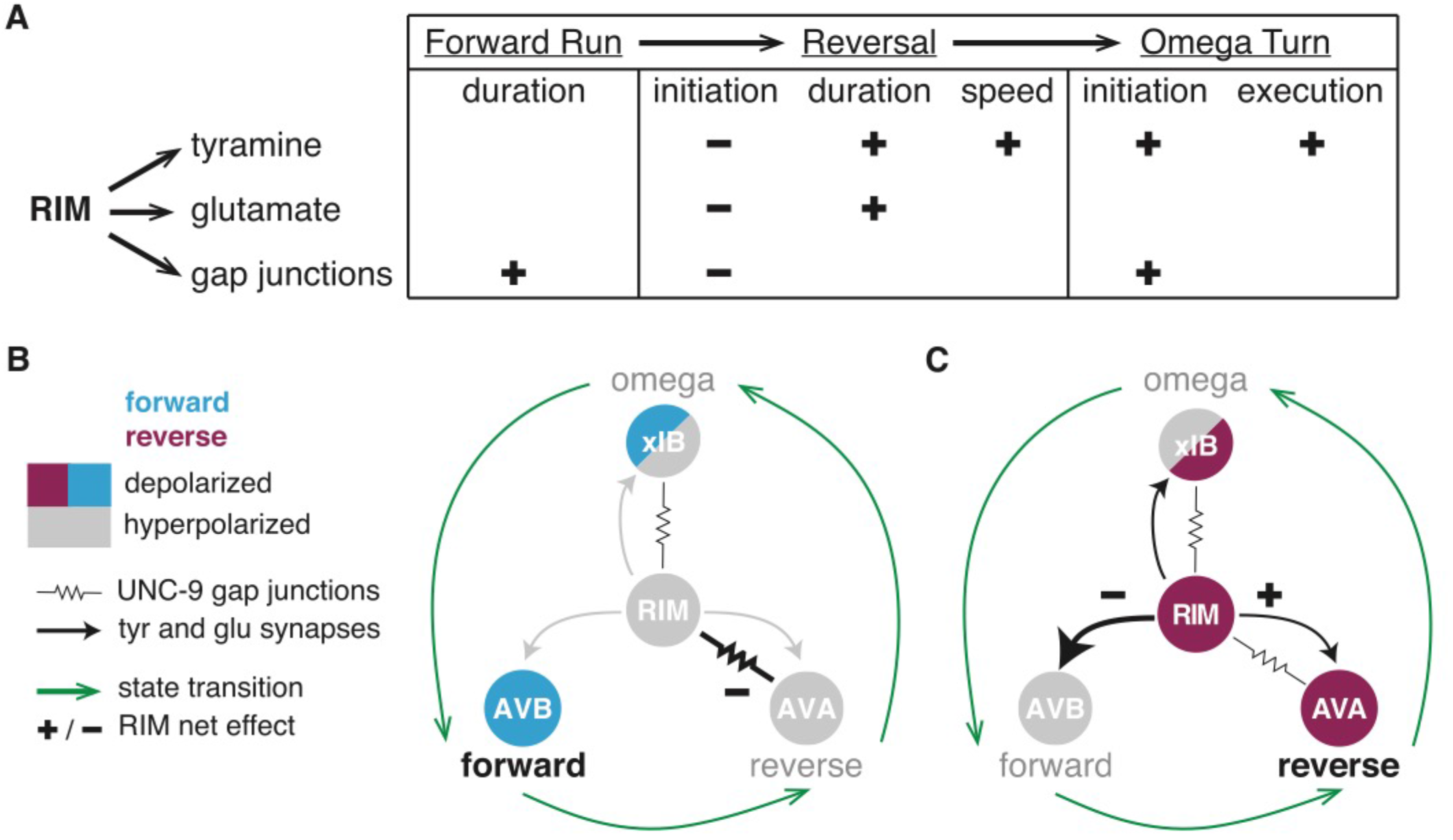
RIM synapses generate behavioral inertia and jointly regulate forward-to-reversal transitions. **(A)** Summary of synaptic regulation of spontaneous behaviors (Figures 2-7). (+) indicates that the normal function of the synapse increases the behavioral parameter (e.g. RIM tyramine increases reversal speed, because *tdc-1* mutant reversals are slower than wild-type). (-) indicates that the synapse decreases the parameter (e.g. RIM tyramine, glutamate, and *unc-1/unc-9* gap junctions all inhibit reversal initiation, because the mutants have more spontaneous reversals than wild-type). Additional RIM transmitters inhibit reversals during global search (Figure 4), and RIC octopamine increases forward locomotion speed (Figure 2–figure supplement 2, Figure 4–figure supplement 1). **(B-C)** AVB, AVA, and xIB (AIB + RIB) are representative of the neurons that promote forward runs, reversals, and omega turns, respectively. AVB and RIB are depolarized during forward runs; RIM, AVA, and AIB are depolarized during reversals; AIB and RIB are depolarized during turns. **(B)** RIM *unc-1/unc-9* gap junctions stabilize forward runs by propagating a hyperpolarizing signal to reversal-promoting neurons. **(C)** RIM tyramine and glutamate stabilize reversals by inhibiting forward-promoting neurons and may also activate reversal-promoting neurons.

### RIM neurotransmitters cooperate to stabilize reversals

RIM controls specific locomotor features: it increases spontaneous reversal speed and duration (Gray et al., 2005, this work), suppresses head oscillations during reversals, and sharpens the omega turns coupled to reversals (Alkema et al., 2005; Donnelly et al., 2013; Pirri et al., 2009), These functions all rely on the RIM transmitter tyramine, which also increases the length of reversals evoked by aversive sensory stimuli (Alkema et al., 2005; Pirri et al., 2009). We found that RIM glutamate increases spontaneous reversal length and duration, but only during the coupled reversal-omega maneuver, and does not increase reversal speed. Both RIM glutamate and tyramine also extend reversals evoked by acute depolarization.

Neurons that release both classical transmitters, like glutamate, and biogenic amines, like tyramine, can employ them additively, cooperatively, or distinctly. In mice, dopaminergic neurons that project from the ventral tegmental area to the nucleus accumbens release both dopamine and glutamate, and either transmitter can support positive reinforcement (Zell et al., 2020). In both *Drosophila* and mice, the glutamate transporter enhances dopamine loading into synaptic vesicles for a cooperative effect (Aguilar et al., 2017; Munster-Wandowski et al., 2016). At a more subtle level, GABA and dopamine co-released from terminals in the mammalian striatum affect target neurons differently – GABA rapidly inhibits action potentials, while dopamine modulates activity through slower GPCR pathways (Tritsch & Sabatini, 2012). The non-additive effects of RIM glutamate and tyramine on spontaneous and optogenetically-evoked behavioral dynamics suggest that they act as co-transmitters to cooperatively stabilize reversal states when RIM is depolarized.

*C. elegans* glutamate receptors and tyramine receptors are broadly expressed in the locomotor circuit. Among RIM’s synaptic targets, the AIB and AVA reversal-promoting neurons express excitatory AMPA-type glutamate receptors, as does RIM itself (Brockie et al., 2001; Hart et al., 1995; Taylor et al., 2019), and glutamate is released onto AVA during reversals (Marvin et al., 2013). RIM glutamate might reinforce the reversal state by depolarizing AVA, while RIM tyramine inhibits the competing forward state via the tyramine-gated chloride channel LGC-55 on AVB (Pirri et al., 2009). In addition, the AVB and RIB forward-promoting neurons express a number of inhibitory glutamate-gated chloride channels (GluCls), and inhibition of AVB could be a site of RIM glutamate-tyramine convergence (Brockie & Maricq, 2006; Dent et al., 2000; Taylor et al., 2019). However, AIB, AVA, and RIM also express GluCls that regulate reversals (Li et al., 2020), and RIM expresses LGC-55 (Taylor et al., 2019). Since all of these receptors co-exist in a circuit rich in positive and negative feedback (Roberts et al., 2016), cell-specific knockouts of receptors as well as neurotransmitters may be needed to define their functions precisely.

### RIM gap junctions stabilize forward runs

For both chemogenetic hyperpolarization and optogenetic depolarization of RIM, the effects on reversal frequency were opposite to those predicted from RIM ablation. Hyperpolarization led to an unanticipated increase in forward run durations, pointing to an active function for RIM when silenced. RIM and its gap junction partners AVA, AVE, and AIB have low activity during forward locomotion; our results suggest that in the hyperpolarized forward state, RIM gap junctions inhibit the AVA backward command neurons and possibly others as well (Gordus et al., 2015; Kagawa-Nagamura et al., 2018; Kato et al., 2015). This hypothesis was supported by the *unc-1(dn)* inhibition of *unc-9*-containing gap junctions in RIM, which shortened the long forward runs induced by RIM hyperpolarization. Moreover, expression of *unc-1(dn)* in the wild-type RIM reduced forward run durations, again, linking these gap junctions in RIM to the hyperpolarized state.

Our conclusion that RIM gap junctions stabilize a hyperpolarized state resonates with previous studies in a different part of the reversal circuit. In addition to their gap junctions with RIM, the AVA neurons form gap junctions with *unc-9*-expressing VA and DA motor neurons that drive backward locomotion. Genetic inactivation of those gap junctions results in defects in forward locomotion and increases calcium levels in AVA (Kawano et al., 2011). From this result, the UNC-9-UNC-7 gap junctions were inferred to decrease the activity of AVA based on hyperpolarizing current flow from VA and DA motor neurons. This role is similar to the role we propose for gap junctions between AVA and RIM. In fact, the *unc-9* innexin expressed in VA/DA neurons and RIM can form heterotypic gap junctions with the *unc-*7 innexin expressed in AVA (Kawano et al., 2011). However, unlike the RIM gap junction knockdown, which acts primarily to affect the duration of coordinated forward runs, the AVA-motor neuron knockdown results in highly uncoordinated movement.

The experiments here, and in Kawano (2011), are limited by the fact that behavior and calcium imaging do not directly measure gap junction conductances. Moreover, direct measurements of gap junctions between AVA and VA5 motor neurons indicate that UNC-9-UNC-7 gap junctions transmit depolarizing current from VA5 to AVA (P. Liu et al., 2017; Shui et al., 2020). That said, reconstitution in *Xenopus* oocytes revealed a startling array of properties among UNC-9-UNC-7 gap junctions, depending on which of seven UNC-7 splice forms is expressed (Shui et al., 2020). How innexins and their splice forms contribute to RIM-to-AVA communication, other than requiring *unc-9* function in RIM, remains to be determined.

Interactions between chemical and electrical synapses play prominent roles in motor circuits including the stomatogastric ganglion of crustaceans, the heartbeat circuit in leeches, and rapid escape circuits in nematodes, arthropods, and fish (Kristan et al., 2005; Marder, 1998; Szczupak, 2016). Although chemical synapses in these circuits can be either excitatory or inhibitory, their electrical synapses have mainly been thought to be excitatory. We speculate that inhibitory electrical synapses resembling those of RIM gap junctions may emerge as stabilizing elements of other motor circuits with long-lasting, mutually exclusive states.

Optogenetic depolarization of RIM elicits reversals, an effect that is reciprocal to that of RIM hyperpolarization. While gap junctions from RIM to AVA could be attractive candidates for this activity, the overall increase in reversal frequency upon RIM depolarization was only slightly diminished by the *unc-9* gap junction knockdown and unaffected by RIM chemical transmitters. RIM expresses 11 innexin genes, the most of any neuron (Bhattacharya et al., 2019). RIM gap junctions may depolarize AVA via innexins that are not affected by the *unc-1(dn)* transgene, such as *inx-1* (Hori et al., 2018; Li et al., 2020; Wang et al., 2020). Cell-specific knockout of *inx-1* and other innexins should provide deeper understanding of RIM gap junctions in AVA, AIB, AVE, and other neurons.

### RIM regulates motor state transitions

The dynamic functions of RIM in spontaneous motor state transitions during local search are regulated by the combined action of tyramine, glutamate and gap junction signaling. All of these synaptic outputs inhibit reversal initiation, even though RIM glutamate and tyramine stabilize the reversal once it has begun.

Among the characterized neurons within the foraging circuit, RIM is the only neuron where ablation has opposite effects on the initiation and execution of a behavioral state (Gray et al., 2005). The dynamic transition from forward to backward locomotion requires coordinated activity changes across the circuit, with positive and negative feedback between forward- and reversal-active neurons (Roberts et al., 2016) (Figure 1A). A role for RIM gap junctions in preventing reversals is consistent with its proposed action in the hyperpolarized (forward) state, but tyramine and glutamate release are likely to rely upon depolarization. In one model, a low level of neurotransmitter release during forward-to-reversal transitions might oppose reversal initiation, while higher levels promote it. Low-level release would be consistent with the graded electrical properties of many *C. elegans* neurons, including motor neurons (Q. Liu et al., 2009) and RIM (Q. Liu et al., 2018), which can also result in graded transmitter release.

We note, however, that chronic or developmental effects of tyramine might also contribute to the increased reversal frequency upon RIM ablation, *tdc-1* mutations, or tetanus toxin expression. Tyramine release during learning can lead to long-term circuit remodeling, and tyramine mediates systemic responses to starvation and other stresses (De Rosa et al., 2019; Ghosh et al., 2016; Jin et al., 2016; Ozbey et al., 2020). Reversal frequencies during local search are regulated by prior experience on bacterial food, including its density and distribution (Calhoun et al., 2014; Lopez-Cruz et al., 2019); tyramine is a candidate to mediate this longer-term behavioral effect.

These transitions, as well as the interactions between RIM, AIB, and RIB that promote transitions from reversals to omega turns, deserve fuller scrutiny (Wang et al., 2020). Here, we have focused on high-resolution quantitative analysis of behavioral parameters, extending the approach of Roberts et al. (2016) and others, to complement the increasingly rich studies of neuronal activity associated with locomotion (Ji et al., 2019; Kato et al., 2015; Kaplan et al., 2020; Nguyen et al., 2016; Venkatachalam et al., 2016). Integration of high-resolution behavior with high-resolution imaging is a critical next step in understanding transition dynamics.

## MATERIALS AND METHODS

### Nematode and Bacterial Culture

In all experiments bacterial food was *E. coli* strain OP50. Nematodes were grown at room temperature (21-22°C) or at 20°C on nematode growth media plates (NGM; 51.3 mM NaCl, 1.7% agar, 0.25% peptone, 1 mM CaCl2, 12.9 μM cholesterol, 1 mM MgSO4, 25 mM KPO4, pH 6) seeded with 200 µL of a saturated *E. coli* liquid culture grown in LB at room temperature or at 37°C, and stored at 4°C (Brenner, 1974). All experiments were performed on young adult hermaphrodites, picked as L4 larvae the evening before an experiment.

Wild-type controls are derived from the N2 Bristol strain, and an additional wild-type strain CX0007 was derived by loss of the transgene from CX17882, to maximize the similarity of controls within an experiment. Mutant strains were backcrossed into N2 at least 3x to reduce background mutations. Wild-type controls in all figures were matched to test strains for transgenes and co-injection markers. Supplementary file 1 includes full genotypes and detailed descriptions of all strains and transgenes.

### Molecular Biology and Transgenics

A 4.5 kilobase region upstream of *tdc-1* that drives expression in RIM and RIC neurons was used for all RIM manipulations. In all cases other than the RIM glutamate knockout, these reagents affect RIC as well as RIM. To separate the functions of the RIM and RIC neurons, we used a 4.5 kilobase region upstream of *tbh-1* to drive expression in RIC neurons. Phenotypes specific to the *tdc-1* transgenes were inferred to have an essential contribution from RIM. See Supplementary file 1 and Supplementary file 2 for relevant strains and plasmids.

Transgenic animals were generated by microinjecting the relevant plasmid of interest with a fluorescent co-injection marker (*myo-2p*::mCherry, *myo-3p*::mCherry, *elt-2p*::nGFP, *elt-2p*::mCherry, *unc-122p*::GFP) and empty pSM vector to reach a final DNA concentration of 100 ng/µL. Transgenes were maintained as extrachromosomal arrays.

### Foraging Assay

Off-food foraging assays were performed and analyzed following López-Cruz et al., 2019. Animals were first preconditioned to a homogenous *E. coli* lawn to standardize their behavioral state (Calhoun et al., 2014). The homogenous lawn was made by seeding NGM plates with a thin uniform layer of saturated *E. coli* liquid culture ∼16 hours before the beginning of the assay. 20 young adult hermaphrodites were placed on this lawn for 45 minutes prior to recording and constrained to a fixed area of 25 cm^2^ using filter paper soaked in 20 mM CuCl_2_. 12-15 of these preconditioned animals were transferred to an unseeded NGM plate briefly to clear adherent bacteria, and then transferred to a large unseeded NGM assay plate, where they were constrained to a fixed area of ∼80 cm^2^ using filter paper soaked in 20 mM CuCl_2_. Video recordings of these animals began within 5-6 minutes from food removal to capture local search behavior. Animals were recorded for 45 minutes as previously described using a 15 MP PL-D7715 CMOS video camera (Pixelink). Frames were acquired at 3 fps using Streampix software (Norpix), using four cameras to image four assays in parallel. LED backlights (Metaphase Technologies) and polarization sheets were used to achieve uniform illumination (López-Cruz et al., 2019). Each experimental strain was assayed a minimum of six times over two days, with control strains assayed in parallel. Animals were tracked using custom MATLAB software (BargmannWormTracker) without manual correction of tracks (López-Cruz et al., 2019, Pokala et al., 2014). Tracker software is available at: https://github.com/navinpokala/BargmannWormTracker.

### Quantification of Spontaneous Behavior

Behavioral states were extracted from the State array generated by BargmannWormTracker. Local search event frequencies per minute were calculated 4-9 min after removal from food. Global search frequencies per minute were calculated 36-41 min after removal from food. Only tracks that were continuous for the entire five-minute time interval were included in frequency analysis. When calculating frequencies, tracks taken on a single day from a single assay plate were averaged to give a single data point, e.g. in Figure 2B and 2D.

Distributions of reversal parameters and forward run durations were calculated using events observed during local search, 4-9 minutes after removal from food. All reversals were included; only forward runs over 2 s in length were included. Reversal length is the path length calculated using the X-Y coordinates, worm length, and pixel size extracted from the tracker. Reversal and forward run speed are the average of mean and median speed extracted from the tracker. Tracks that were less than 3 minutes long, tracks approaching the copper barrier, and tracks that did not include a complete reversal or forward run were not included in reversal and forward run parameter analyses.

### Optogenetic Manipulations

The red-shifted channelrhodopsin ReaChR (Lin et al., 2013) was expressed under the *tdc-1* promoter and animals were stimulated during the off-food foraging assay described above, following López-Cruz et al., 2019. Experimental animals were treated with 12.5 µM all-trans retinal overnight and during preconditioning on homogeneous food lawns; control animals were prepared in parallel on plates that did not contain retinal. Optogenetic stimuli were delivered with a 525 nm Solis HighPower LED (ThorLabs) controlled by custom MATLAB software and strobed at a 5% duty cycle. 2 (Figures 5, Figure 6–figure supplement 1) or 3 (Figures 8, Figure 7–figure supplement 2) pulses of ∼45 µW/mm^2^ light delivered for 20 s each with a 100-s interpulse interval starting at 8 or 10 min (local search) and 38 or 40 min (global search). These light intensities elicited the maximal behavioral effect of ReaChR. Additional lower light intensities (not shown) were included in each experiment, with pulses always separated by 100 s.

For behavioral quantification, tracks were aligned around the light pulses and extracted over a 120 s period, with the light pulse delivered at 50-70 s. Only tracks that were continuous for the entire 120 s period were used. The change in reversal frequency was calculated by subtracting the mean reversal frequency during an 18-s time window before light onset from the mean reversal frequency during an 18-s time window during the light pulse. Behavioral parameters were scored only for the first reversal of duration ≥0.5 s that began during the light stimulation.

### Histamine Treatment

The *Drosophila* histamine-gated chloride channel HisCl1 was expressed under the *tdc-1* promoter. Animals were treated with histamine following Pokala et al., 2014. Histamine dihydrochloride (Sigma-Aldrich H7250) was dissolved in Milli-Q® purified water, filtered, and stored at −20°C. Histamine solution was added to cooled NGM (45-50°C) for a final concentration of 10 mM to make assay plates. Animals were habituated and cleaned on NGM plates that did not contain histamine and then recorded on 10 mM histamine assay plates for 45 minutes. See Quantification of Spontaneous Behavior above.

### Statistical Analyses

All statistical analyses were conducted in GraphPad Prism except for the two-sample Kolmogorov Smirnov test, which was performed in MATLAB. When making multiple comparisons, the p-values of the two-sample Kolmogorov Smirnov test were adjusted with a Bonferroni correction. The effect size was calculated for all significant distribution comparisons as the D statistic, which represents the maximum distance between the empirical cumulative distributions of the data. Because of the large n values in these experiments, even very small effects reached statistical significance. Based on control strains (e.g. Figure 2–figure supplement 1), we set a meaningful effect size of ≥0.15 as a cutoff for interpreting results, See Supplementary file 3 for the n of all distributions. See Supplementary file 4 for a summary of all p-values and statistical tests.

## ACKNOWLEDGEMENTS

We thank Philip Kidd, Andrew Gordus, Qiang Liu, Elias Scheer, Javier Marquina-Solis, Audrey Harnagel, James Lee, Friederike Buck, Leslie Vosshall, Vanessa Ruta, Yishi Jin, and Jeremy Dittman for thoughtful discussions and comments on the manuscript. We thank Alejandro López-Cruz for his collaboration in creating the cell-specific *eat-4* knockout strain. This work was supported by the Howard Hughes Medical Institute, of which CIB was an investigator, and by the Chan Zuckerberg Initiative.

---

**Figure 1–figure supplement 1.**
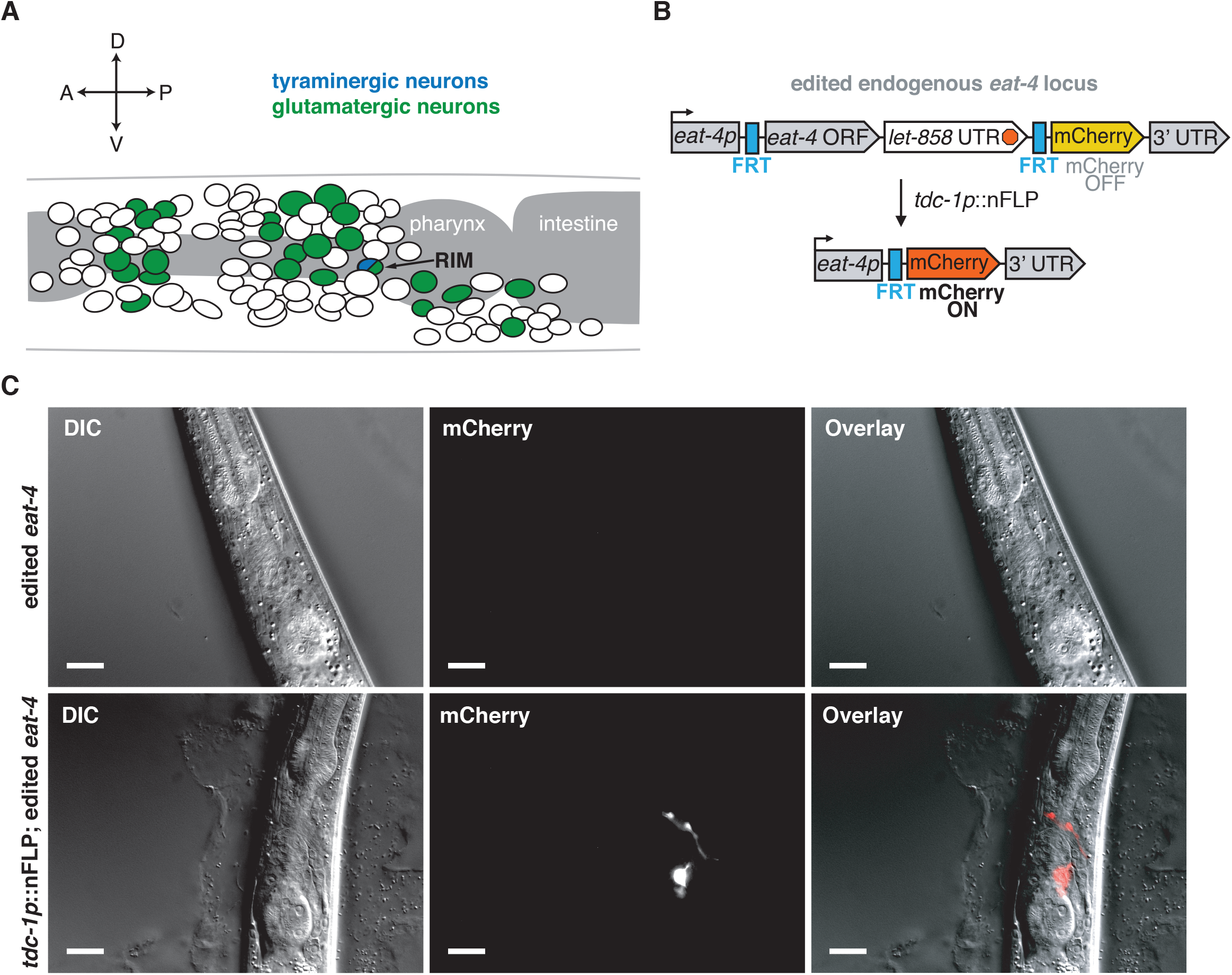
CRISPR-Cas9 generated alleles enable RIM-specific glutamate transporter knockout. **(A)** Schematic of the sources of glutamate and tyramine in the *C. elegans* nervous system. Adapted from Pereira et al., 2015. **(B)** Schematic of cell-specific glutamate transporter knockout genetic strategy. Using CRISPR-Cas9, an FRT site was inserted immediately before the start codon of *eat-4* (VGLUT) and *let-858* 3’-UTR::FRT::mCherry immediately after the stop codon of *eat-4*. *let-858* 3’-UTR stops transcription so mCherry is not expressed. To knock out glutamate release in this edited strain, nuclear-localized flippase (nFLP) was expressed under a *tdc-1* promoter. The intersection of *tdc-1* and *eat-4* expression is limited to RIM, leading to excision of the *eat-4* ORF in RIM, confirmed by mCherry expression in the targeted cells. **(C)** Validation of CRISPR-Cas9 recombination. Top panel: Differential interference contrast (DIC) (left), mCherry fluorescence (middle), and merged (right) images of animals with edited endogenous *eat-4* (VGLUT) locus. In the edited strain there is no mCherry expression, confirming that the *let-858* 3’-UTR stops transcription. Bottom: DIC (left), mCherry (middle), and merged (right) images for edited *eat-4* strain following RIM-specific nFLP expression (*tdc-1*p::nFLP). mCherry is expressed only in RIM neurons, the intersection of *eat-4* and *tdc-1* expression. mCherry specificity for RIM was confirmed by screening 30 animals. Scale bar is 20 μM.

**Figure 2–figure supplement 1.**
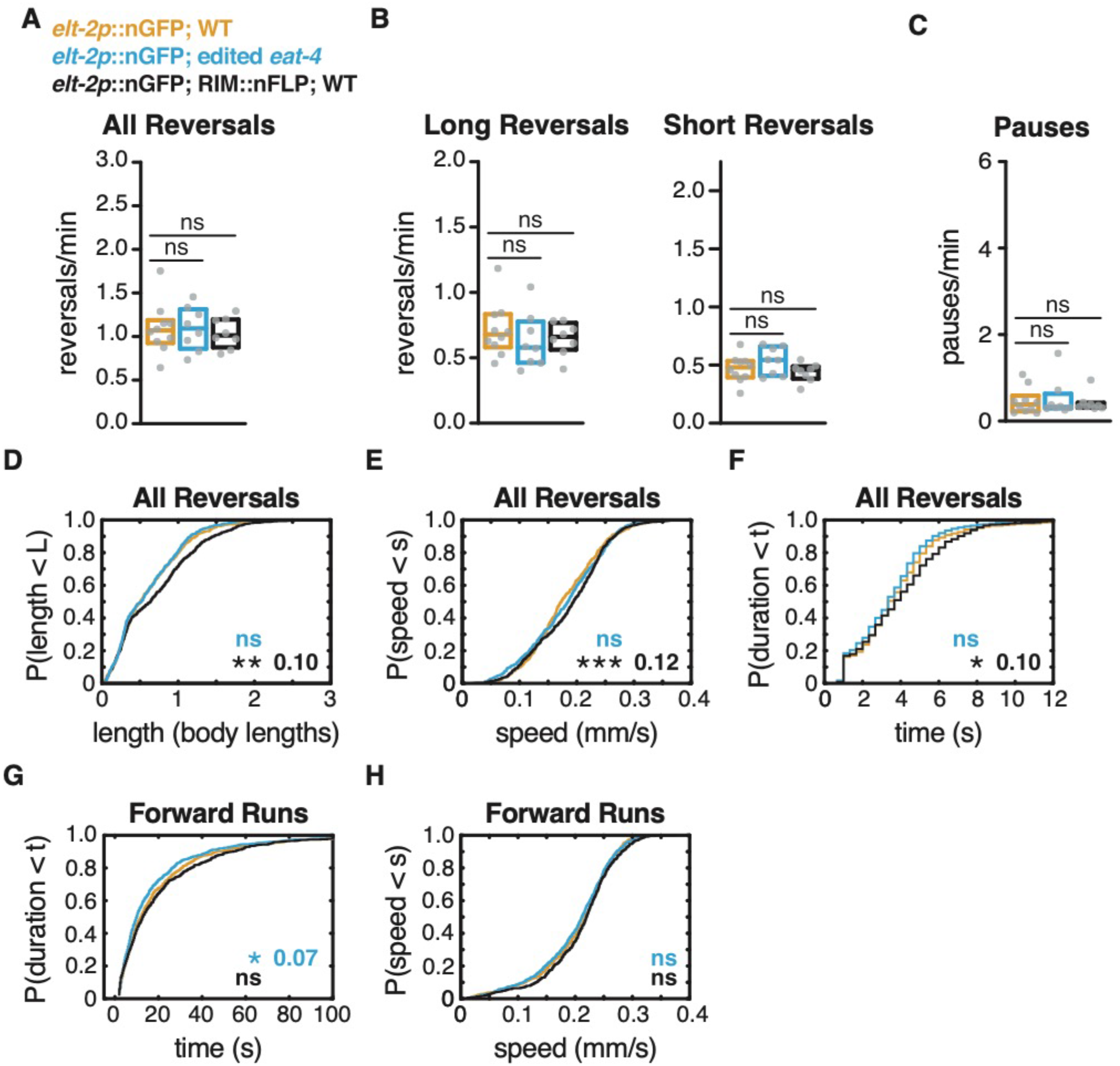
RIM::nFLP and edited *eat-4* do not account for RIM glu KO phenotype. **(A)** Mean frequency of all reversals during local search (4-9 min off food). **(B)** Mean frequency of long reversals (>0.5 body lengths, left) and short reversals (<0.5 body lengths, right) during local search. **(C)** Mean frequency of pauses during local search. **(D-F)** For all reversals during local search, empirical cumulative distributions of reversal length **(D)** reversal speed **(E)** and reversal duration **(F)**. n = 584-642 reversals per genotype from 8-10 assays, 12-15 animals per assay (Supplementary file 3). **(G-H)** For all forward runs ≥2 s in duration during local search, empirical cumulative distributions of run duration **(G)** and run speed **(H).** n = 687-913 runs per genotype from 8-10 assays, 12-15 animals per assay (Supplementary file 3). **(A-C)** Each gray dot is the mean for 12-15 animals on a single assay plate, with 22-24 plates per genotype. Boxes indicate median and interquartile range for all assays. Asterisks indicate statistical significance compared to *elt-2p*::nGFP; WT using a Mann-Whitney test (ns = p-value >= 0.05). **(D-H)** Asterisks indicate statistical significance compared to WT using a two-sample Kolmogorov Smirnov test (* = p-value < 0.05, *** = p-value < 0.0001, ns = p-value >= 0.05). Numbers in figures indicate effect size (See Materials and methods).

**Figure 2–figure supplement 2.**
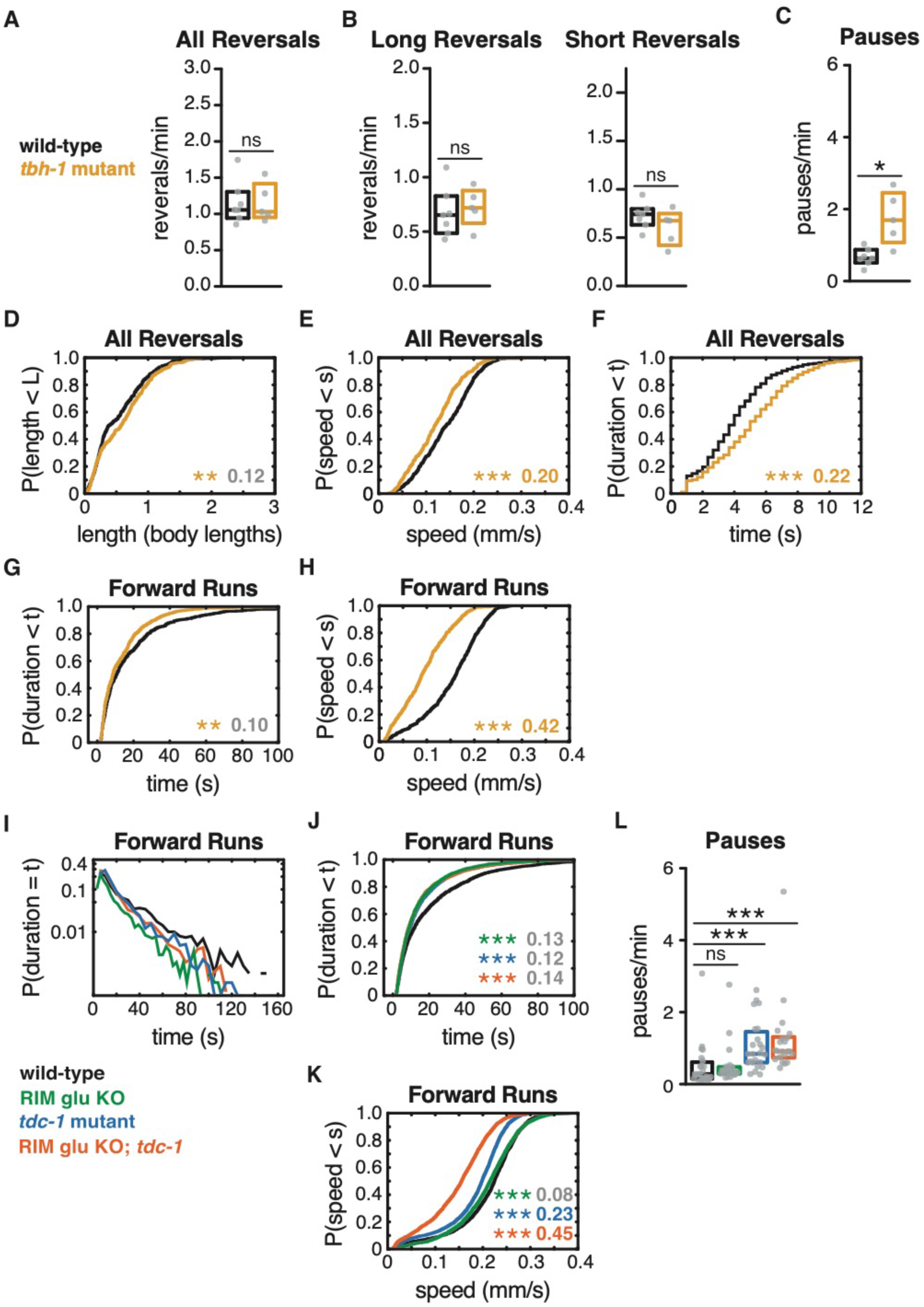
Octopamine affects forward and reversal speed but not reversal length or frequency. **(A)** Mean frequency of all reversals during local search (4-9 min off food). **(B)** Mean frequency of long reversals (>0.5 body lengths, left) and short reversals (<0.5 body lengths, right) during local search. **(C)** Mean frequency of pauses during local search. **(D-F)** For all reversals during local search, empirical cumulative distributions of reversal length **(D)** reversal speed **(E)** and reversal duration **(F)**. n = 384-532 reversals per genotype from 5-7 assays, 12-15 animals per assay (Supplementary file 3). **(G-H)** For all forward runs ≥2 s in duration during local search, empirical cumulative distributions of run duration **(G)** and run speed **(H).** n = 618-636 runs per genotype from 5-7 assays, 12-15 animals per assay (Supplementary file 3). **(I)** Forward run durations for genotypes in Figures 1-3 follow an exponential distribution. Y-axis set at a log10 scale. **(J-K)** For all forward runs ≥2 s in duration during local search for genotypes in Figures 1-3, empirical cumulative distributions of run duration **(J)** and run speed **(K)**. n = 1898-3132 runs per genotype from 22-24 assays, 12-15 animals per assay (Supplementary file 3). **(L)** Mean frequency of pauses during local search for genotypes in Figures 1-3. **(A-C, L)** Each gray dot is the mean for 12-15 animals on a single assay plate, with 5-7 **(A-C)** or 22-24 **(L)** plates per genotype. Boxes indicate median and interquartile range for all assays. Asterisks indicate statistical significance compared to WT using a Mann-Whitney **(A-C)** or Kruskal-Wallis test with Dunn’s multiple comparisons test **(L)** (* = p-value < 0.05, *** = p-value < 0.0001, ns = p-value >= 0.05). **(D-H, J-K)** Asterisks indicate statistical significance compared to WT using a two-sample Kolmogorov Smirnov test with **(J,K)** or without **(D-H)** a Bonferroni correction. (* = p-value < 0.05, *** = p-value < 0.0001, ns = p-value >= 0.05). Numbers in figures indicate effect size (See Materials and methods). Although statistically significant, the effect sizes indicated in gray in D, G, J, and K fell below the 0.15 cutoff for interpretation established using control strains. The *tbh-1* mutant **(H)** accounts for the effect of *tdc-1* **(K)** on forward speed. *tbh-1* mutants, which are deficient in the synthesis of octopamine, were deficient in reversal speed to a lesser extent than *tdc-1* mutants, without decreasing reversal length. We conclude that tyramine increases reversal length and speed. *tbh-1* mutants had a large reduction in forward speed, comparable to that of *tdc-1* mutants, and also increased pausing rates and decreased forward run duration.

**Figure 4–figure supplement 1.**
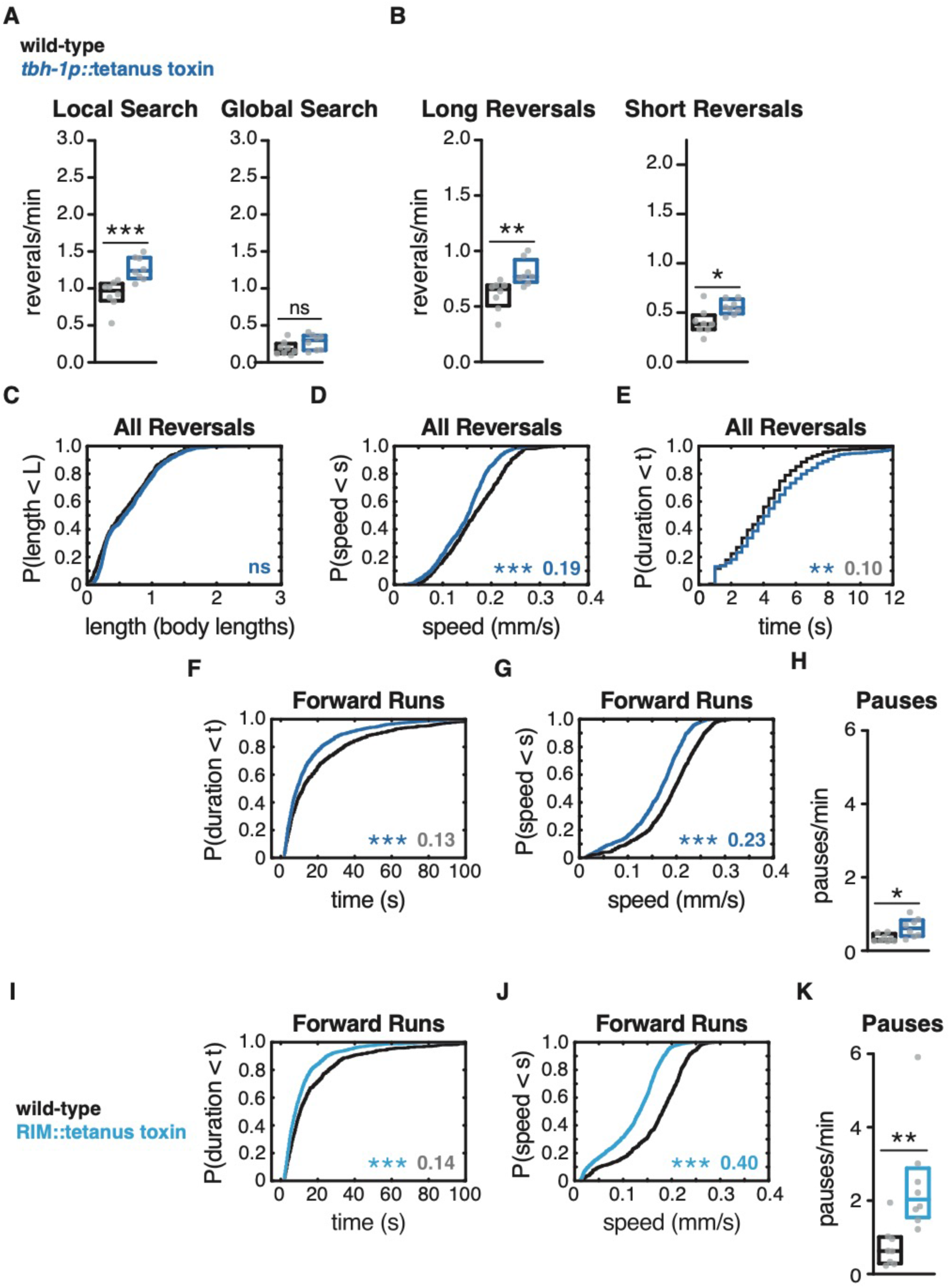
Expression of tetanus toxin in RIC does not affect reversal length or global search reversal frequency. Mean frequency of all reversals during local search (4-9 min off food, left) and global search (36-41 min off food, right). **(A)** Mean frequency of long reversals (>0.5 body lengths, left) and short reversals (<0.5 body lengths, right) during local search. **(C-E)** For all reversals during local search, empirical cumulative distributions of reversal length **(B)** reversal speed **(D)** and reversal duration **(E)**. n = 504-733 reversals per genotype from 8 assays, 12-15 animals per assay (Supplementary file 3). **(F-G, I-J)** For all forward runs ≥2 s in duration during local search, empirical cumulative distributions of run duration **(F, I)** and run speed **(G, J)** in animals expressing tetanus toxin in RIC **(F-G)** or RIM + RIC **(I-J).** n = 657-1301 runs per genotype from 8 assays, 12-15 animals per assay (Supplementary file 3). **(H, K)** Mean frequency of pauses during local search in animals expressing tetanus toxin in RIC **(H)** or RIM + RIC **(K)**. **(A-C, H, K)** Each gray dot is the mean for 12-15 animals on a single assay plate, with 8 plates per genotype. Boxes indicate median and interquartile range for all assays. Asterisks indicate statistical significance compared to WT using a Mann-Whitney test (* = p-value < 0.05, ** = p-value < 0.01, *** = p-value < 0.001, ns = p-value >= 0.05). **(C-G, I-J)** Asterisks indicate statistical significance compared to WT using a two-sample Kolmogorov Smirnov test. (** = p-value < 0.01, *** = p-value < 0.0001, ns = p-value >= 0.05). Numbers in figures indicate effect size. Although statistically significant, the effect sizes indicated in gray in E, F, and I fell below the 0.15 cutoff for interpretation established using control strains. Expressing tetanus toxin in RIC caused a small increase in both long and short reversals during local search and had small effects on reversal parameters. Note that the small effects on reversal parameters suggest that RIC is not the critical source of tyramine in Figure 3. RIC tetanus toxin had effects similar to *tbh-1* mutants on forward and reversal speed and pausing. We conclude that RIM tetanus toxin (Figure 4) has significant effects on reversal frequency, length, speed, and duration, in local and global search.

**Figure 4–figure supplement 2.**
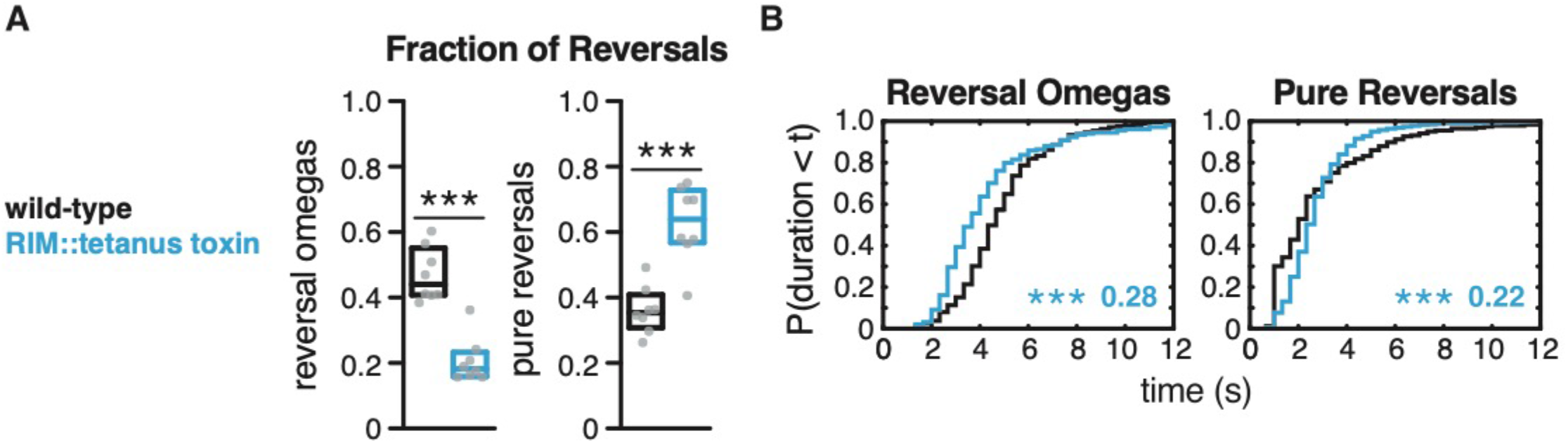
Expression of tetanus toxin in RIM decreases reversal-omega coupling and alters reversal omega and pure reversal duration. **(A)** Fraction of reversals that terminate in an omega turn (left) and fraction of pure reversals (right) for each genotype (see Figure 2F-G). Each gray dot is the mean for 12-15 animals on a single assay plate, with 8 plates per genotype. Boxes indicate median and interquartile range for all assays. Asterisks indicate statistical significance compared to WT using a Mann-Whitney test (*** = p-value < 0.001). **(B)** For reversal-omega maneuvers (left) and pure reversals (right) during local search, empirical cumulative distributions of reversal durations. Asterisks indicate statistical significance compared to WT using a two-sample Kolmogorov Smirnov test (*** = p-value < 0.0001). Numbers in figures indicate effect size. n = 211-647 events per genotype from 8 assays, 12-15 animals per assay (Supplementary file 3).

**Figure 6–figure supplement 1.**
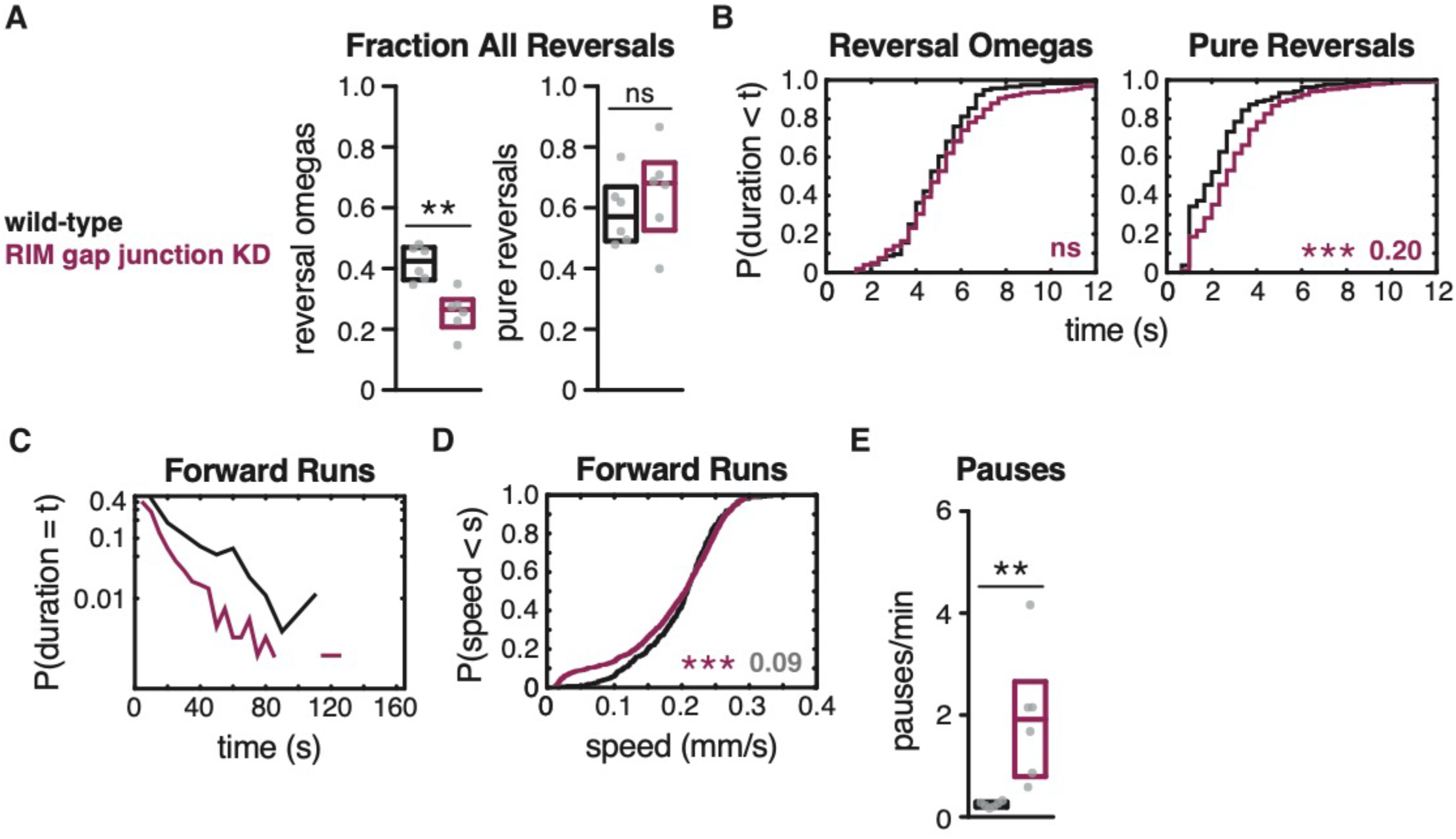
RIM gap junctions affect reversal-omega and pausing frequency. **(A)** Fraction of reversals that terminate in an omega turn (left) and fraction of pure reversals (right) for each genotype (see Figure 2F-G). **(B)** For reversal-omega maneuvers (left) and pure reversals (right) during local search, empirical cumulative distributions of reversal durations. n = 116-473 events per genotype from 6 assays, 12-15 animals per assay (Supplementary file 3). **(C)** Exponential plot of forward run durations. Y-axis set at a log10 scale (see Figure 6H). **(D)** For all forward runs ≥2 s in duration during local search, empirical cumulative distributions of run speed. n = 355-889 runs per genotype from 6 assays, 12-15 animals per assay (Supplementary file 3). **(E)** Mean frequency of pauses during local search. **(A, E)** Each gray dot is the mean for 12-15 animals on a single assay plate, with 6 plates per genotype. Boxes indicate median and interquartile range for all assays. Asterisks indicate statistical significance compared to WT using a Mann-Whitney test (** = p-value < 0.01, ns = p-value >= 0.05). **(B, D)** Asterisks indicate statistical significance compared to WT using a two-sample Kolmogorov Smirnov (*** = p-value < 0.001, ns = p-value >= 0.05). Numbers in figures indicate effect size. Effect sizes indicated in gray fall below the 0.15 cutoff for interpretation established using control strains. RIM gap junction KD decreased the fraction of reversal-omegas without altering reversal-omega durations.

**Figure 6–figure supplement 2.**
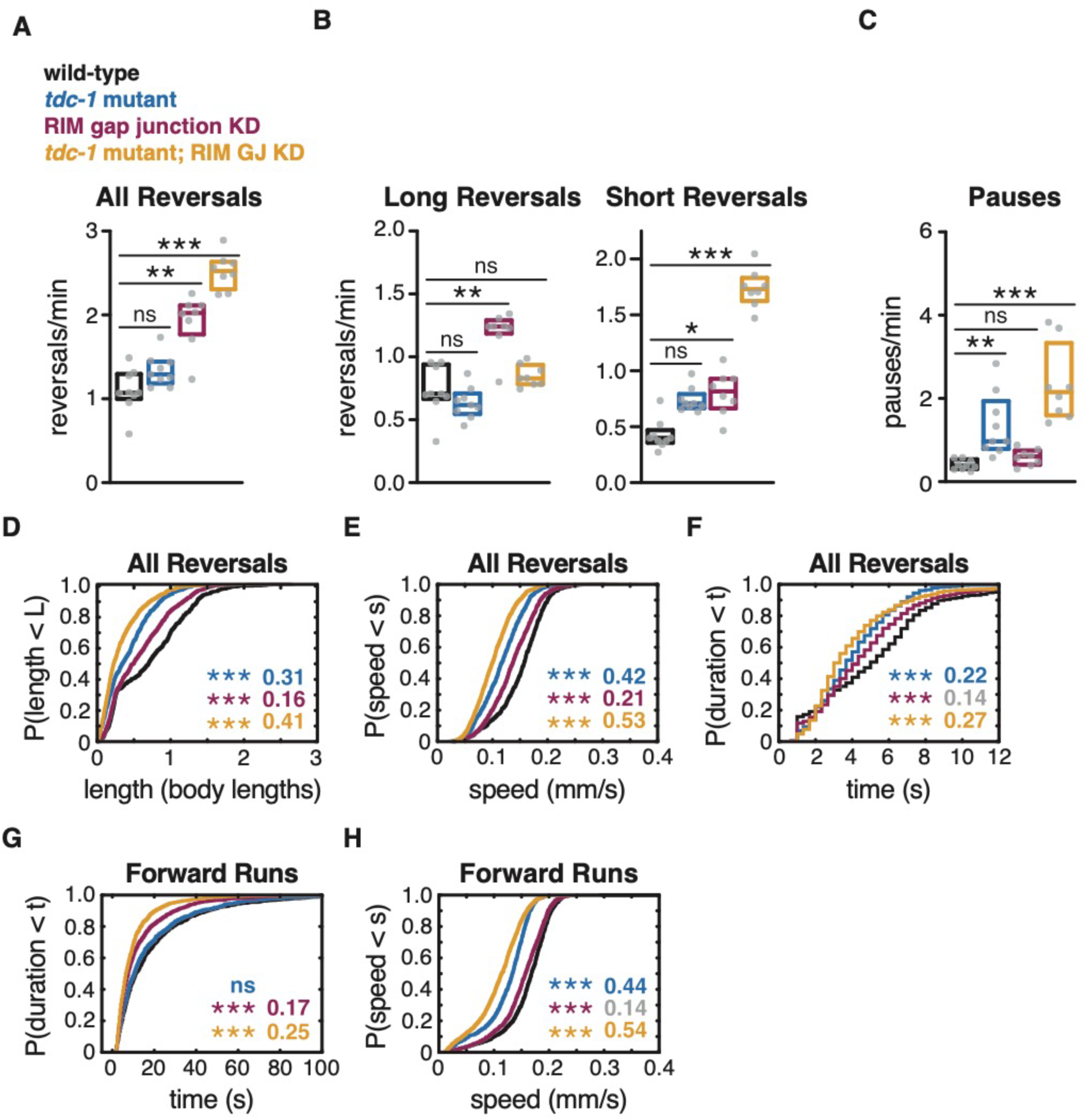
RIM gap junctions and RIM tyramine act additively in spontaneous local search behavior. **(A)** Mean frequency of all reversals during local search (4-9 min off food). **(B)** Mean frequency of long reversals (>0.5 body lengths, left) and short reversals (<0.5 body lengths, right) during local search. **(C)** Mean frequency of pauses during local search. **(D-F)** For all reversals during local search, empirical cumulative distributions of reversal length **(D)** reversal speed **(E)** and reversal duration **(F)**. n = 691-1432 reversals per genotype from 8-9 assays, 12-15 animals per assay (Supplementary file 3). **(G-H)** For all forward runs ≥2 s in duration during local search, empirical cumulative distributions of run duration **(G)** and run speed **(H).** n = 875-1541 runs per genotype from 8-9 assays, 12-15 animals per assay (Supplementary file 3). **(A-C)** Each gray dot is the mean for 12-15 animals on a single assay plate, with 8-9 plates per genotype. Boxes indicate median and interquartile range for all assays. Asterisks indicate statistical significance compared to WT using a Kruskal-Wallis test with Dunn’s multiple comparisons test (* = p-value < 0.05, ** = p-value < 0.01, *** = p-value < 0.001, ns = p-value ≥ 0.05). **(D-H)** Asterisks indicate statistical significance compared to WT using a two-sample Kolmogorov Smirnov test with a Bonferroni correction (*** = p-value < 0.0001, ns = p-value >= 0.05). Numbers in figures indicate effect size. Effect sizes indicated in gray fall below the 0.15 cutoff for interpretation established using control strains.

**Figure 7–figure supplement 1.**
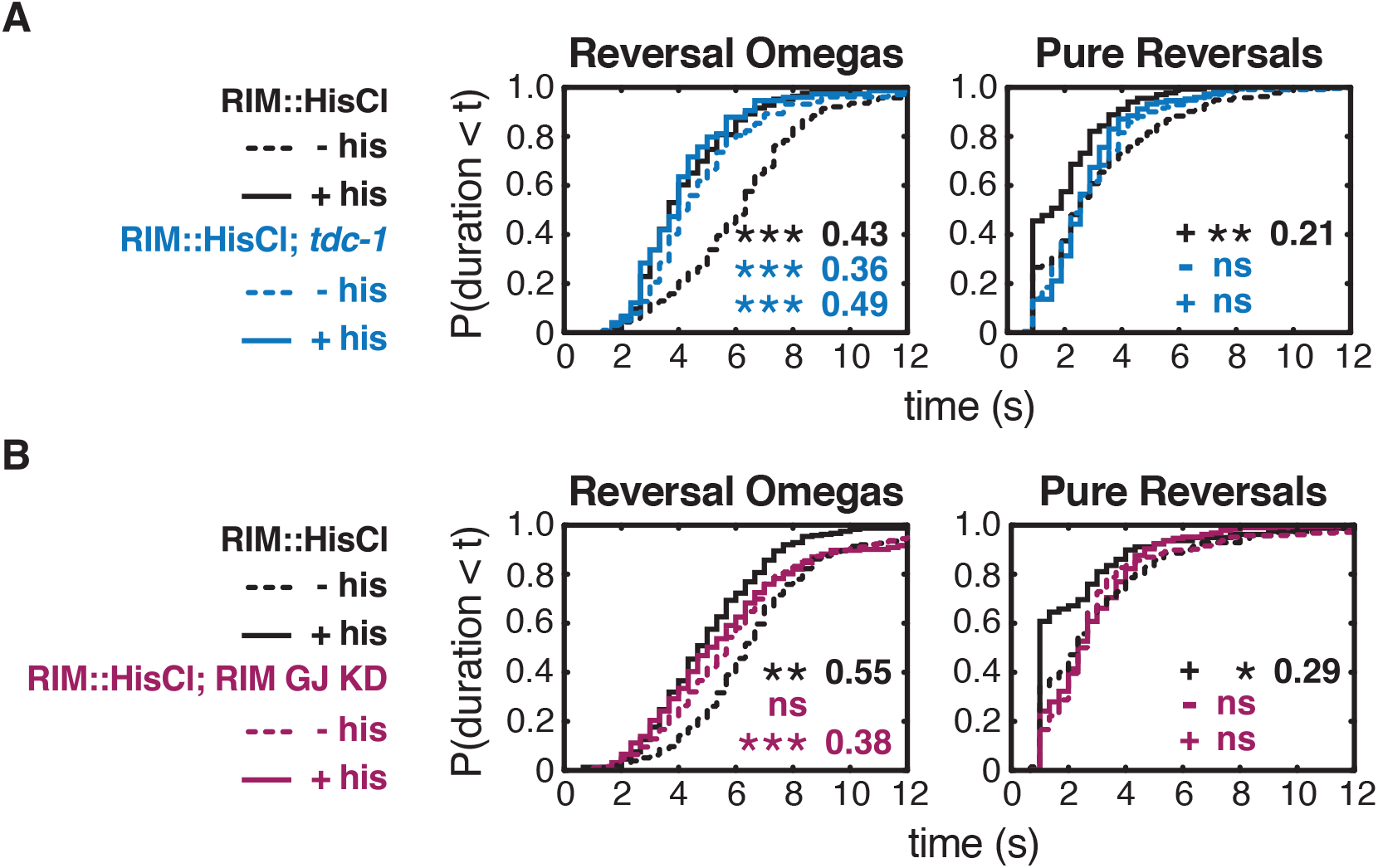
RIM hyperpolarization affects pure reversals via tyramine and gap junctions. **(A,B)** For reversal-omega maneuvers (left) and pure reversals (right) during local search, empirical cumulative distributions of reversal durations in RIM::HisCl strains, including controls **(A,B)**, *tdc-1* **(A)**, and RIM gap junction knockdown **(B),** with and without histamine. **(A)** n = 74-223 events per genotype from 6-8 assays, 12-15 animals per assay (Supplementary file 3). **(B)** n = 15-306 events per genotype from 6-8 assays, 12-15 animals per assay (Supplementary file 3). Hyperpolarizing RIM in a wild-type background resulted in pure reversals of ≤1 s that were not normally observed during spontaneous behavior. These very short reversals were suppressed by *tdc-1* mutation and gap junction knockdown.

**Figure 7–figure supplement 2.**
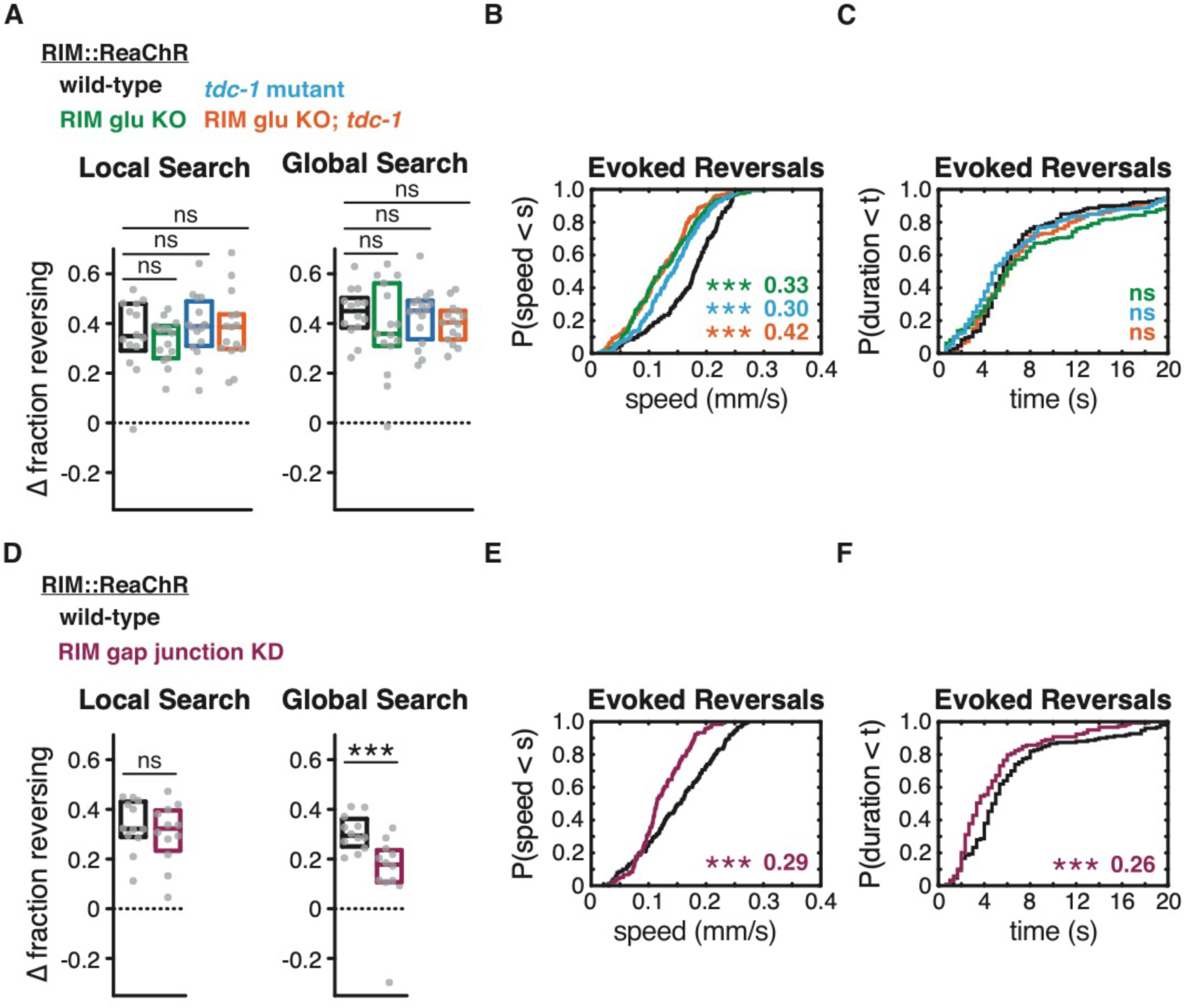
RIM gap junctions support optogenetically-evoked reversals. **(A-F)** Effects of RIM::ReaChR activation in wild-type, RIM glu KO, *tdc-1* mutants, RIM glu KO; *tdc-1* double mutants, and RIM gap junction knockdown animals. All animals pre-treated with all-trans retinal. **(A, D)** Average increase in the fraction of animals reversing during the light pulse during local search (8-14 min off food, left) or global search (38-44 min off food, right). Each gray dot is the mean for 12-15 animals on a single assay plate. Boxes indicate median and interquartile range for all assays. Asterisks indicate statistical significance compared to WT using a Kruskal-Wallis test with Dunn’s multiple comparisons test **(A)** or Mann-Whitney test **(D)** (*** = p-value < 0.001, ns = p-value >= 0.05). **(B-C, E-F)** For all reversals induced during the light pulse during local search (8-14 min), empirical cumulative distributions of reversal speed **(B, E)** and reversal duration **(C, F)**. **(B-C)** n = 159-193 reversals from 14-15 assays, 12-15 animals per assay, two light pulses per assay, 10-14 minutes after removal from food (Supplementary file 3). Asterisks indicate statistical significance compared to controls of the same genotype using a two-sample Kolmogorov Smirnov test with a Bonferroni correction (** = p-value < 0.01, *** = p-value < 0.001). Numbers indicate effect size. **(E-F)** n = 119-150 reversals from 12 assays, 12-15 animals per assay, three light pulses per assay, 8-14 minutes after removal from food (Supplementary file 3). Asterisks indicate statistical significance compared to controls of the same genotype using a two-sample Kolmogorov Smirnov test (** = p-value < 0.01, *** = p-value < 0.001). Numbers indicate effect size. RIM gap junction knockdown, but not RIM chemical synapse mutants, suppressed optogenetically-evoked reversals during global search. RIM gap junction knockdown also decreased optogenetically-evoked reversal duration. Both RIM gap junction knockdown and chemical synapse mutants decreased optogenetically-evoked reversal speed.

