## Supplemental Tables for "Behavioral control by depolarized and hyperpolarized states of an integrating neuron by"

**Supplementary File 1: Table S1. Strain Details**

| **Figures** | **Strain** | **Strain Name** | **Genotype** | **Notes** |
| --- | --- | --- | --- | --- |
| 1-3,  2-S1,  2-S2 | WT | CX17882 | *kyEx6329*[*tdc-1p*::nFLP, *elt-2p*::nGFP] | parent of CX0007 |
| 1-3,  2-S2 | RIM glu KO | CX17881 | *eat-4(ky1091) eat-4(ky1092)* *III; kyEx6329*[*tdc-1p*::nFLP, *elt-2p*::nGFP] | *eat-4(ky1091) eat-4(ky1092)* backcrossed 3x to N2 See López-Cruz et al., 2019 |
| 1-3,  2-S2 | *tdc-1* | CX17883 | *tdc-1(n3419)* *II; kyEx6329*[*tdc-1p*::nFLP, *elt-2p*::nGFP] | *tdc-1(n3419)* *II* backcrossed 11x to N2 |
| 1-3,  2-S2 | RIM glu KO; *tdc-1* | CX17884 | *tdc-1(n3419)* *II;* *eat-4(ky1091) eat-4(ky1092)* *III; kyEx6329*[*tdc-1p*::nFLP, *elt-2p*::nGFP] | *tdc-1(n3419)* *II* backcrossed 11x to N2 *eat-4(ky1091) eat-4(ky1092)* backcrossed 3x to N2 See López-Cruz et al., 2019 |
| 2-S1 | *elt-2p*::nGFP; WT | CX18118 | *kyEx6458*[*elt-2p*::nGFP] | wild-type (CX0007) co-injection marker control; original strain used in experiments lost |
| 2-S1 | *elt-2p*::nGFP; edited *eat-4* | CX17461 x  CX18118 | *eat-4(ky1091) eat-4(ky1092)* *III; kyEx6458*[*elt-2p*::nGFP] | co-injection marker & *eat-4(ky1091) eat-4(ky1092)* control; original strain used in experiments lost |
| 2-S2 | *tbh-1* | MT9455 | *tbh-1(n3247)* *X* | backcrossed to N2 8x for autosomes, 4x for *X* |
| 4, 4-S1, 4-S2,  6-S2 | WT | N2 | N2 | 6-S2 animals derived from transgene negative CX14853 |
| 4, 4-S1, 4-S2 | RIM::tetanus toxin | CX14993 | *kyEx4962*[*tdc-1p*::tetanus toxin::mCherry] |  |
| 4-S1 | *tbh-1p*::tetanus toxin | CX17912 | *kyEx6345*[*tbh-1p*::tetanus toxin::mCherry, *unc-1*22*p*::GFP] |  |
| 5, 7,  7-S1 | RIM::HisCl; WT | CX18193 | *kyIs779*[*tdc-1p*::HisCl::SL2::mCherry] | *kyIs779* = apparent spontaneous integrant of *kyEx5464*; not backcrossed parent: CX16040 |
| 5, 7-S1 | RIM::HisCl; *tdc-1* | CX18194 | *tdc-1(n3419)* *II ;* *kyIs779*[*tdc-1p*::HisCl::SL2::mCherry] | *tdc-1(n3419)* *II* backcrossed 11x to N2  *kyIs779* = apparent spontaneous integrant of *kyEx5464*; not backcrossed  parent: CX16040 |
| 6, 4-S1, 6-S1 | WT | CX17546 | *kyEx6164*[*unc-1*22*p*::GFP] |  |
| 6, 6-S1, 6-S2 | RIM gap junction KD | CX14853 | *kyEx487*[*tdc-1p*::*unc-1(n494)*::SL2::mCherry, *unc-1*22*p*::GFP] |  |
| 6-S2 | *tdc-1* | MT13113 | *tdc-1(n3419)* *II* |  |
| 6-S2 | RIM gap junction KD; *tdc-1* | CX14853 X  MT13113 | *tdc-1(n3419)* *II; kyEx4871*[*tdc-1p*::*unc-1(n494)*::SL2::mCherry, *unc-1*22*p*::GFP] |  |
| 7, 7-S1 | RIM::HisCl; RIM gap junction KD | CX18137 | *kyIs779*[*tdc-1p*::HisCl::SL2::mCherry]; *kyEx6465*[*tdc-1p*::*unc-1(n494)*::SL2::GFP, *elt-2p*::mCherry] | *kyIs779* = apparent spontaneous integrant of *kyEx5464;* not backcrossed parent: CX16040 |
| 7, 7-S2 | RIM::ReaChR: wild-type | CX17885 | *kyEx6329*[*tdc-1p*::nFLP, *elt-2p*::nGFP]; *kyEx6232*[*tdc-1p*::ReaChR::SL2::GFP, *myo-3p*::mCherry] |  |
| 7, 7-S2 | RIM::ReaChR: RIM glu KO | CX17886 | *eat-4(ky1091) eat-4(ky1092)* *III; kyEx6329*[*tdc-1p*::nFLP, *elt-2p*::nGFP]; *kyEx6232*[*tdc-1p*::ReaChR::SL2::GFP, *myo-3p*::mCherry] |  |
| 7, 7-S2 | RIM::ReaChR: *tdc-1* | CX17887 | *tdc-1(n3419) II;*  *kyEx6329*[*tdc-1p*::nFLP, *elt-2p*::nGFP]; *kyEx6232*[*tdc-1p*::ReaChR::SL2::GFP, *myo-3p*::mCherry] | *tdc-1(n3419)* *II* backcrossed 11x to N2 |
| 7, 7-S2 | RIM::ReaChR: RIM glu KO; *tdc-1* | CX17888 | *tdc-1(n3419)* *II;* *eat-4(ky1091) eat-4(ky1092)* *III; kyEx6329*[*tdc-1p*::nFLP, *elt-2p*::nGFP] *kyEx6232*[*tdc-1p*::ReaChR::SL2::GFP, *myo-3p*::mCherry] | *tdc-1(n3419)* *II* backcrossed 11x to N2 *eat-4*(*ky1091) eat-4*(*ky1092)* backcrossed 3x to N2 See López-Cruz et al., 2019 |
| 7, 7-S2 | RIM::ReaChR: wild-type | CX17694 | *kyEx6232*[*tdc-1p*::ReaChR::SL2::GFP, *myo-3p*::mCherry] |  |
| 7, 7-S2 | RIM::ReaChR: RIM gap junction KD | CX18195 | *kyEx6232*[*tdc-1p*::ReaChR::SL2::GFP, *myo-3p*::mCherry]; *kyEx4871*[*tdc-1p*::*unc-1(n494)*::SL2::mCherry, *unc-1*22*p*::GFP] |  |

**Supplementary File 2: Table S2. Plasmids Generated for This Study**

| **Name** | **Description** | **Figures** |
| --- | --- | --- |
| pAS03 | *tdc-1p*::nFLP | 1-3, 7, S2, S3, S9 |
| pAS04 | *tbh-1p*::tetanus toxin::mCherry | S4 |
| pAS05 | *tdc-1p*::*unc-1(n494)*::SL2::GFP | 7, S8 |

**Supplementary File 3: Table S3. Reversals and Forward Runs, n values**

| **Figures** | **Strain** | **All Reversals** | **Reversal Omegas** | **Pure Reversals** | **Forward Runs** |
| --- | --- | --- | --- | --- | --- |
| 2-3, 2-S2 | WT | 1443 | 605 | 555 | 1898 |
| 2-3, 2-S2 | RIM glu KO | 2689 | 1142 | 1011 | 3121 |
| 2-3, 2-S2 | *tdc-1* | 2760 | 628 | 1462 | 3132 |
| 2-3, 2-S2 | RIM glu KO; *tdc-1* | 2217 | 500 | 1231 | 2594 |
| 2-S1 | *elt-2p*::nGFP; WT | 642 |  |  | 861 |
| 2-S1 | *elt-2p*::nGFP; edited *eat-4* | 634 |  |  | 913 |
| 2-S1 | *elt-2p*::nGFP; tdc-1*p*::nFLP; WT | 584 |  |  | 687 |
| 2-S2 | wild-type | 532 |  |  | 636 |
| 2-S2 | *tbh-1* | 384 |  |  | 618 |
| 4, 4-S1, 4-S2 | WT | 595 | 239 | 211 | 657 |
| 4, 4-S1, 4-S2 | RIM::tetanus toxin | 1066 | 185 | 647 | 1301 |
| 4-S1 | WT | 504 |  |  | 664 |
| 4-S1 | tbh-1*p*::tetanus toxin | 733 |  |  | 1041 |
| 5, 7-S1 | RIM::HisCl; WT (- his) | 503 | 139 | 223 | 579 |
| 5, 7-S1 | RIM::HisCl; WT (+ his) | 259 | 83 | 138 | 399 |
| 5, 7-S1 | RIM::HisCl; *tdc-1* (- his) | 440 | 102 | 218 | 545 |
| 5, 7-S1 | RIM::HisCl; *tdc-1* (+ his) | 251 | 74 | 148 | 566 |
| 6, 6-S1 | WT | 330 | 116 | 134 | 355 |
| 6, 6-S1 | RIM gap junction KD | 933 | 193 | 473 | 889 |
| 6-S2 | WT | 691 | *321* | *240* | 875 |
| 6-S2 | *tdc-1* | 813 | *238* | *342* | 1015 |
| 6-S2 | RIM gap junction KD | 1100 | *391* | *473* | 1055 |
| 6-S2 | RIM gap junction KD; *tdc-1* | 1432 | *261* | *909* | 1541 |
| 7, 7-S1 | RIM::HisCl; WT (- his) | 192 | 71 | 70 | 330 |
| 7, 7-S1 | RIM::HisCl; WT (+ his) | 126 | 15 | 79 | 400 |
| 7, 7-S1 | RIM::HisCl; RIM gap junction KD (- his) | 668 | 176 | 306 | 819 |
| 7, 7-S1 | RIM::HisCl; RIM gap junction KD (+ his) | 320 | 69 | 186 | 692 |
| 7, 7-S2 | RIM::ReaChR; wild-type | 167 | *108* | *42* | N/A |
| 7, 7-S2 | RIM::ReaChR; RIM glu KO | 159 | *92* | *54* | N/A |
| 7, 7-S2 | RIM::ReaChR; *tdc-1* | 159 | *84* | *54* | N/A |
| 7, 7-S2 | RIM::ReaChR; RIM glu KO; *tdc-1* | 193 | *122* | *55* | N/A |
| 7, 7-S2 | RIM::ReaChR; WT | 150 | *96* | *48* | N/A |
| 7, 7-S2 | RIM::ReaChR; RIM gap junction KD | 119 | *54* | *58* | N/A |
| Italics designate n not included in manuscript. | | | | | |

**Supplementary File 4: Table S4. Statistical Analyses**

| **Panel** | **Comparison** | **Summary** | **p-value** | **alpha** | **Test** |
| --- | --- | --- | --- | --- | --- |
| 2B; left | WT vs RIM glu KO | *** | <0.0001 | 0.05 | Kruskal-Wallis; Dunn's correction |
| 2B; left | WT vs *tdc-1* | *** | <0.0001 | 0.05 | Kruskal-Wallis; Dunn's correction |
| 2B; left | WT vs RIM glu KO; *tdc-1* | *** | 0.0050 | 0.05 | Kruskal-Wallis; Dunn's correction |
| 2B; right | WT vs RIM glu KO | ns | >0.9999 | 0.05 | Kruskal-Wallis; Dunn's correction |
| 2B; right | WT vs *tdc-1* | ns | 0.4410 | 0.05 | Kruskal-Wallis; Dunn's correction |
| 2B; right | WT vs RIM glu KO; *tdc-1* | ns | 0.0614 | 0.05 | Kruskal-Wallis; Dunn's correction |
| 2D; left | WT vs RIM glu KO | *** | <0.0001 | 0.05 | Kruskal-Wallis; Dunn's correction |
| 2D; left | WT vs *tdc-1* | ns | >0.9999 | 0.05 | Kruskal-Wallis; Dunn's correction |
| 2D; left | WT vs RIM glu KO; *tdc-1* | ns | >0.9999 | 0.05 | Kruskal-Wallis; Dunn's correction |
| 2D; right | WT vs RIM glu KO | *** | 0.0003 | 0.05 | Kruskal-Wallis; Dunn's correction |
| 2D; right | WT vs *tdc-1* | *** | <0.0001 | 0.05 | Kruskal-Wallis; Dunn's correction |
| 2D; right | WT vs RIM glu KO; *tdc-1* | *** | <0.0001 | 0.05 | Kruskal-Wallis; Dunn's correction |
| 2G; left | WT vs RIM glu KO | ns | 0.9869 | 0.05 | Kruskal-Wallis; Dunn's correction |
| 2G; left | WT vs *tdc-1* | *** | <0.0001 | 0.05 | Kruskal-Wallis; Dunn's correction |
| 2G; left | WT vs RIM glu KO; *tdc-1* | *** | <0.0001 | 0.05 | Kruskal-Wallis; Dunn's correction |
| 2G; right | WT vs RIM glu KO | ns | >0.9999 | 0.05 | Kruskal-Wallis; Dunn's correction |
| 2G; right | WT vs *tdc-1* | ** | 0.0056 | 0.05 | Kruskal-Wallis; Dunn's correction |
| 2G; right | WT vs RIM glu KO; *tdc-1* | *** | <0.0001 | 0.05 | Kruskal-Wallis; Dunn's correction |
| 2H | WT vs RIM glu KO | ns | >0.9999 | 0.05 | Kruskal-Wallis; Dunn's correction |
| 2H | WT vs *tdc-1* | *** | <0.0001 | 0.05 | Kruskal-Wallis; Dunn's correction |
| 2H | WT vs RIM glu KO; *tdc-1* | *** | <0.0001 | 0.05 | Kruskal-Wallis; Dunn's correction |
| 2-S1A | *elt-2p*::nGFP; WT vs *elt-2p*::nGFP; edited *eat-4* | ns | 0.9470 | 0.05 | Mann-Whitney |
| 2-S1A | *elt-2p*::nGFP; WT vs *elt-2p*::nGFP; *tdc-1p*::nFLP; WT | ns | 0.8785 | 0.05 | Mann-Whitney |
| 2-S1B; left | *elt-2p*::nGFP; WT vs *elt-2p*::nGFP; edited *eat-4* | ns | 0.3154 | 0.05 | Mann-Whitney |
| 2-S1B; left | *elt-2p*::nGFP; WT vs *elt-2p*::nGFP; *tdc-1p*::nFLP; WT | ns | 0.6965 | 0.05 | Mann-Whitney |
| 2-S1B; right | *elt-2p*::nGFP; WT vs *elt-2p*::nGFP; edited *eat-4* | ns | 0.2112 | 0.05 | Mann-Whitney |
| 2-S1B; right | *elt-2p*::nGFP; WT vs *elt-2p*::nGFP; *tdc-1p*::nFLP; WT | ns | 0.5726 | 0.05 | Mann-Whitney |
| 2-S1C | *elt-2p*::nGFP; WT vs *elt-2p*::nGFP; edited *eat-4* | ns | 0.8460 | 0.05 | Mann-Whitney |
| 2-S1C | *elt-2p*::nGFP; WT vs *elt-2p*::nGFP; *tdc-1p*::nFLP; WT | ns | 0.9473 | 0.05 | Mann-Whitney |
| 2-S1D | *elt-2p*::nGFP; WT vs *elt-2p*::nGFP; edited *eat-4* | ns | 0.3153 | 0.05 | 2-sample Kolmogorov-Smirnov |
| 2-S1D | *elt-2p*::nGFP; WT vs *elt-2p*::nGFP; *tdc-1p*::nFLP; WT | ** | 0.0023 | 0.05 | 2-sample Kolmogorov-Smirnov |
| 2-S1E | *elt-2p*::nGFP; WT vs *elt-2p*::nGFP; edited *eat-4* | ns | 0.1197 | 0.05 | 2-sample Kolmogorov-Smirnov |
| 2-S1E | *elt-2p*::nGFP; WT vs *elt-2p*::nGFP; *tdc-1p*::nFLP; WT | *** | 0.0002 | 0.05 | 2-sample Kolmogorov-Smirnov |
| 2-S1F | *elt-2p*::nGFP; WT vs *elt-2p*::nGFP; edited *eat-4* | ns | 0.2853 | 0.05 | 2-sample Kolmogorov-Smirnov |
| 2-S1F | *elt-2p*::nGFP; WT vs *elt-2p*::nGFP; *tdc-1p*::nFLP; WT | ** | 0.0057 | 0.05 | 2-sample Kolmogorov-Smirnov |
| 2-S1G | *elt-2p*::nGFP; WT vs *elt-2p*::nGFP; edited *eat-4* | * | 0.0246 | 0.05 | 2-sample Kolmogorov-Smirnov |
| 2-S1G | *elt-2p*::nGFP; WT vs *elt-2p*::nGFP; *tdc-1p*::nFLP; WT | ns | 0.2849 | 0.05 | 2-sample Kolmogorov-Smirnov |
| 2-S1H | *elt-2p*::nGFP; WT vs *elt-2p*::nGFP; edited *eat-4* | ns | 0.2050 | 0.05 | 2-sample Kolmogorov-Smirnov |
| 2-S1H | *elt-2p*::nGFP; WT vs *elt-2p*::nGFP; *tdc-1p*::nFLP; WT | ns | 0.3725 | 0.05 | 2-sample Kolmogorov-Smirnov |
| 2-S2A | WT vs *tbh-1* | ns | 0.8763 | 0.05 | Mann-Whitney |
| 2-S2B; left | WT vs *tbh-1* | ns | 0.5303 | 0.05 | Mann-Whitney |
| 2-S2B; right | WT vs *tbh-1* | ns | 0.2677 | 0.05 | Mann-Whitney |
| 2-S2C | WT vs *tbh-1* | * | 0.0101 | 0.05 | Mann-Whitney |
| 2-S2D | WT vs *tbh-1* | ** | 0.0046 | 0.05 | 2-sample Kolmogorov-Smirnov |
| 2-S2E | WT vs *tbh-1* | *** | <0.0001 | 0.05 | 2-sample Kolmogorov-Smirnov |
| 2-S2F | WT vs *tbh-1* | *** | <0.0001 | 0.05 | 2-sample Kolmogorov-Smirnov |
| 2-S2G | WT vs *tbh-1* | ** | 0.0019 | 0.05 | 2-sample Kolmogorov-Smirnov |
| 2-S2H | WT vs *tbh-1* | *** | <0.0001 | 0.05 | 2-sample Kolmogorov-Smirnov |
| 2-S2J | WT vs RIM glu KO | *** | <0.0001 | 0.05 | 2-sample Kolmogorov-Smirnov; Bonferroni correction |
| 2-S2J | WT vs *tdc-1* | *** | <0.0001 | 0.05 | 2-sample Kolmogorov-Smirnov; Bonferroni correction |
| 2-S2J | WT vs RIM glu KO; *tdc-1* | *** | <0.0001 | 0.05 | 2-sample Kolmogorov-Smirnov; Bonferroni correction |
| 2-S2K | WT vs RIM glu KO | *** | <0.0001 | 0.05 | 2-sample Kolmogorov-Smirnov; Bonferroni correction |
| 2-S2K | WT vs *tdc-1* | *** | <0.0001 | 0.05 | 2-sample Kolmogorov-Smirnov; Bonferroni correction |
| 2-S2K | WT vs RIM glu KO; *tdc-1* | *** | <0.0001 | 0.05 | 2-sample Kolmogorov-Smirnov; Bonferroni correction |
| 2-S2L | WT vs RIM glu KO | ns | >0.9999 | 0.05 | Kruskal-Wallis; Dunn's correction |
| 2-S2L | WT vs *tdc-1* | *** | 0.0002 | 0.05 | Kruskal-Wallis; Dunn's correction |
| 2-S2L | WT vs RIM glu KO; *tdc-1* | *** | <0.0001 | 0.05 | Kruskal-Wallis; Dunn's correction |
| 3A | WT vs RIM glu KO | *** | <0.0001 | 0.05 | 2-sample Kolmogorov-Smirnov; Bonferroni correction |
| 3A | WT vs *tdc-1* | *** | <0.0001 | 0.05 | 2-sample Kolmogorov-Smirnov; Bonferroni correction |
| 3A | WT vs RIM glu KO; *tdc-1* | *** | <0.0001 | 0.05 | 2-sample Kolmogorov-Smirnov; Bonferroni correction |
| 3B | WT vs RIM glu KO | *** | <0.0001 | 0.05 | 2-sample Kolmogorov-Smirnov; Bonferroni correction |
| 3B | WT vs *tdc-1* | *** | <0.0001 | 0.05 | 2-sample Kolmogorov-Smirnov; Bonferroni correction |
| 3B | WT vs RIM glu KO; *tdc-1* | *** | <0.0001 | 0.05 | 2-sample Kolmogorov-Smirnov; Bonferroni correction |
| 3C | WT vs RIM glu KO | *** | <0.0001 | 0.05 | 2-sample Kolmogorov-Smirnov; Bonferroni correction |
| 3C | WT vs *tdc-1* | *** | <0.0001 | 0.05 | 2-sample Kolmogorov-Smirnov; Bonferroni correction |
| 3C | WT vs RIM glu KO; *tdc-1* | *** | <0.0001 | 0.05 | 2-sample Kolmogorov-Smirnov; Bonferroni correction |
| 3D | WT vs RIM glu KO | *** | <0.0001 | 0.05 | 2-sample Kolmogorov-Smirnov; Bonferroni correction |
| 3D | WT vs *tdc-1* | *** | <0.0001 | 0.05 | 2-sample Kolmogorov-Smirnov; Bonferroni correction |
| 3D | WT vs RIM glu KO; *tdc-1* | *** | <0.0001 | 0.05 | 2-sample Kolmogorov-Smirnov; Bonferroni correction |
| 3E | WT vs RIM glu KO | *** | <0.0001 | 0.05 | 2-sample Kolmogorov-Smirnov; Bonferroni correction |
| 3E | WT vs *tdc-1* | *** | <0.0001 | 0.05 | 2-sample Kolmogorov-Smirnov; Bonferroni correction |
| 3E | WT vs RIM glu KO; *tdc-1* | *** | <0.0001 | 0.05 | 2-sample Kolmogorov-Smirnov; Bonferroni correction |
| 3F | WT vs RIM glu KO | *** | <0.0001 | 0.05 | 2-sample Kolmogorov-Smirnov; Bonferroni correction |
| 3F | WT vs *tdc-1* | *** | <0.0001 | 0.05 | 2-sample Kolmogorov-Smirnov; Bonferroni correction |
| 3F | WT vs RIM glu KO; *tdc-1* | *** | <0.0001 | 0.05 | 2-sample Kolmogorov-Smirnov; Bonferroni correction |
| 3G | WT vs RIM glu KO | * | 0.0444 | 0.05 | 2-sample Kolmogorov-Smirnov; Bonferroni correction |
| 3G | WT vs *tdc-1* | *** | <0.0001 | 0.05 | 2-sample Kolmogorov-Smirnov; Bonferroni correction |
| 3G | WT vs RIM glu KO; *tdc-1* | *** | <0.0001 | 0.05 | 2-sample Kolmogorov-Smirnov; Bonferroni correction |
| 3H | WT vs RIM glu KO | ns | 0.2622 | 0.05 | 2-sample Kolmogorov-Smirnov; Bonferroni correction |
| 3H | WT vs *tdc-1* | *** | <0.0001 | 0.05 | 2-sample Kolmogorov-Smirnov; Bonferroni correction |
| 3H | WT vs RIM glu KO; *tdc-1* | *** | <0.0001 | 0.05 | 2-sample Kolmogorov-Smirnov; Bonferroni correction |
| 3I | WT vs RIM glu KO | ns | 0.8643 | 0.05 | 2-sample Kolmogorov-Smirnov; Bonferroni correction |
| 3I | WT vs *tdc-1* | *** | <0.0001 | 0.05 | 2-sample Kolmogorov-Smirnov; Bonferroni correction |
| 3I | WT vs RIM glu KO; *tdc-1* | *** | <0.0001 | 0.05 | 2-sample Kolmogorov-Smirnov; Bonferroni correction |
| 4C; left | WT vs RIM::tetanus toxin | ** | 0.0070 | 0.05 | Mann-Whitney |
| 4C; right | WT vs RIM::tetanus toxin | * | 0.0281 | 0.05 | Mann-Whitney |
| 4D; left | WT vs RIM::tetanus toxin | ns | 0.1105 | 0.05 | Mann-Whitney |
| 4D; right | WT vs RIM::tetanus toxin | *** | 0.0002 | 0.05 | Mann-Whitney |
| 4E | WT vs RIM::tetanus toxin | *** | <0.0001 | 0.05 | 2-sample Kolmogorov-Smirnov |
| 4F | WT vs RIM::tetanus toxin | *** | <0.0001 | 0.05 | 2-sample Kolmogorov-Smirnov |
| 4G | WT vs RIM::tetanus toxin | *** | <0.0001 | 0.05 | 2-sample Kolmogorov-Smirnov |
| 4-S1A; left | WT vs *tbh-1p*::tetanus toxin | *** | 0.0006 | 0.05 | Mann-Whitney |
| 4-S1A; right | WT vs *tbh-1p*::tetanus toxin | ns | 0.1304 | 0.05 | Mann-Whitney |
| 4-S1B; left | WT vs *tbh-1p*::tetanus toxin | ** | 0.0022 | 0.05 | Mann-Whitney |
| 4-S1B; right | WT vs *tbh-1p*::tetanus toxin | * | 0.0207 | 0.05 | Mann-Whitney |
| 4-S1C | WT vs *tbh-1p*::tetanus toxin | ns | 0.6943 | 0.05 | 2-sample Kolmogorov-Smirnov |
| 4-S1D | WT vs *tbh-1p*::tetanus toxin | *** | <0.0001 | 0.05 | 2-sample Kolmogorov-Smirnov |
| 4-S1E | WT vs *tbh-1p*::tetanus toxin | ** | 0.0055 | 0.05 | 2-sample Kolmogorov-Smirnov |
| 4-S1F | WT vs *tbh-1p*::tetanus toxin | *** | <0.0001 | 0.05 | 2-sample Kolmogorov-Smirnov |
| 4-S1G | WT vs *tbh-1p*::tetanus toxin | *** | <0.0001 | 0.05 | 2-sample Kolmogorov-Smirnov |
| 4-S1H | WT vs *tbh-1p*::tetanus toxin | * | 0.0101 | 0.05 | Mann-Whitney |
| 4-S1I | WT vs RIM::tetanus toxin | *** | <0.0001 | 0.05 | 2-sample Kolmogorov-Smirnov |
| 4-S1J | WT vs RIM::tetanus toxin | *** | <0.0001 | 0.05 | 2-sample Kolmogorov-Smirnov |
| 4-S1K | WT vs RIM::tetanus toxin | ** | 0.0019 | 0.05 | Kruskal-Wallis; Dunn's correction |
| 4-S2A; left | WT vs RIM::tetanus toxin | *** | 0.0002 | 0.05 | Mann-Whitney |
| 4-S2A; right | WT vs RIM::tetanus toxin | *** | 0.0006 | 0.05 | Mann-Whitney |
| 4-S2B; left | WT vs RIM::tetanus toxin | *** | <0.0001 | 0.05 | 2-sample Kolmogorov-Smirnov |
| 4-S2B; right | WT vs RIM::tetanus toxin | *** | <0.0001 | 0.05 | 2-sample Kolmogorov-Smirnov |
| 5C | RIM::HisCl: WT (- his) vs WT (+ his) | ** | 0.0085 | 0.05 | Kruskal-Wallis; Dunn's correction |
| 5C | RIM::HisCl: WT (- his) vs *tdc-1* (- his) | ns | >0.9999 | 0.05 | Kruskal-Wallis; Dunn's correction |
| 5C | RIM::HisCl: WT (- his) vs *tdc-1* (+ his) | ** | 0.0036 | 0.05 | Kruskal-Wallis; Dunn's correction |
| 5D | RIM::HisCl: WT (- his) vs WT (+ his) | *** | <0.0001 | 0.05 | 2-sample Kolmogorov-Smirnov; Bonferroni correction |
| 5D | RIM::HisCl: WT (- his) vs *tdc-1* (- his) | ns | 0.5367 | 0.05 | 2-sample Kolmogorov-Smirnov; Bonferroni correction |
| 5D | RIM::HisCl: WT (- his) vs *tdc-1* (+ his) | *** | <0.0001 | 0.05 | 2-sample Kolmogorov-Smirnov; Bonferroni correction |
| 5E | RIM::HisCl: WT (- his) vs WT (+ his) | *** | <0.0001 | 0.05 | 2-sample Kolmogorov-Smirnov; Bonferroni correction |
| 5E | RIM::HisCl: WT (- his) vs *tdc-1* (- his) | *** | <0.0001 | 0.05 | 2-sample Kolmogorov-Smirnov; Bonferroni correction |
| 5E | RIM::HisCl: WT (- his) vs *tdc-1* (+ his) | *** | <0.0001 | 0.05 | 2-sample Kolmogorov-Smirnov; Bonferroni correction |
| 5F | RIM::HisCl: WT (- his) vs WT (+ his) | *** | <0.0001 | 0.05 | 2-sample Kolmogorov-Smirnov; Bonferroni correction |
| 5F | RIM::HisCl: WT (- his) vs *tdc-1* (- his) | *** | <0.0001 | 0.05 | 2-sample Kolmogorov-Smirnov; Bonferroni correction |
| 5F | RIM::HisCl: WT (- his) vs *tdc-1* (+ his) | *** | <0.0001 | 0.05 | 2-sample Kolmogorov-Smirnov; Bonferroni correction |
| 5G | RIM::HisCl: WT (- his) vs WT (+ his) | *** | <0.0001 | 0.05 | 2-sample Kolmogorov-Smirnov; Bonferroni correction |
| 5G | RIM::HisCl: WT (- his) vs *tdc-1* (- his) | *** | <0.0001 | 0.05 | 2-sample Kolmogorov-Smirnov; Bonferroni correction |
| 5G | RIM::HisCl: WT (- his) vs *tdc-1* (+ his) | *** | <0.0001 | 0.05 | 2-sample Kolmogorov-Smirnov; Bonferroni correction |
| 6C; left | WT vs RIM gap junction KD | ** | 0.0022 | 0.05 | Mann-Whitney |
| 6C; right | WT vs RIM gap junction KD | * | 0.0411 | 0.05 | Mann-Whitney |
| 6D; left | WT vs RIM gap junction KD | ** | 0.0022 | 0.05 | Mann-Whitney |
| 6D; right | WT vs RIM gap junction KD | ** | 0.0022 | 0.05 | Mann-Whitney |
| 6E | WT vs RIM gap junction KD | ** | 0.0058 | 0.05 | 2-sample Kolmogorov-Smirnov |
| 6F | WT vs RIM gap junction KD | *** | 0.0008 | 0.05 | 2-sample Kolmogorov-Smirnov |
| 6G | WT vs RIM gap junction KD | ns | 0.4306 | 0.05 | 2-sample Kolmogorov-Smirnov |
| 6H | WT vs RIM gap junction KD | *** | <0.0001 | 0.05 | 2-sample Kolmogorov-Smirnov |
| 6-S1A; left | WT vs RIM gap junction KD | ** | 0.0043 | 0.05 | Mann-Whitney |
| 6-S1A; right | WT vs RIM gap junction KD | ns | 0.3939 | 0.05 | Mann-Whitney |
| 6-S1B; left | WT vs RIM gap junction KD | ns | 0.2682 | 0.05 | 2-sample Kolmogorov-Smirnov |
| 6-S1B; right | WT vs RIM gap junction KD | *** | 0.0004 | 0.05 | 2-sample Kolmogorov-Smirnov |
| 6-S1D | WT vs RIM gap junction KD | *** | <0.0001 | 0.05 | 2-sample Kolmogorov-Smirnov |
| 6-S1E | WT vs RIM gap junction KD | ** | 0.0022 | 0.05 | Mann-Whitney |
| 6-S2A | WT vs *tdc-1* | ns | 0.6683 | 0.05 | Kruskal-Wallis; Dunn's correction |
| 6-S2A | WT vs RIM gap junction KD | ** | 0.0099 | 0.05 | Kruskal-Wallis; Dunn's correction |
| 6-S2A | WT vs *tdc-1*; RIM gap junction KD | *** | <0.0001 | 0.05 | Kruskal-Wallis; Dunn's correction |
| 6-S2B; left | WT vs *tdc-1* | ns | 0.5666 | 0.05 | Kruskal-Wallis; Dunn's correction |
| 6-S2B; left | WT vs RIM gap junction KD | ** | 0.0052 | 0.05 | Kruskal-Wallis; Dunn's correction |
| 6-S2B; left | WT vs *tdc-1*; RIM gap junction KD | ns | 0.7123 | 0.05 | Kruskal-Wallis; Dunn's correction |
| 6-S2B; right | WT vs *tdc-1* | ns | 0.0993 | 0.05 | Kruskal-Wallis; Dunn's correction |
| 6-S2B; right | WT vs RIM gap junction KD | * | 0.0250 | 0.05 | Kruskal-Wallis; Dunn's correction |
| 6-S2B; right | WT vs *tdc-1*; RIM gap junction KD | *** | <0.0001 | 0.05 | Kruskal-Wallis; Dunn's correction |
| 6-S2C | WT vs *tdc-1* | *** | <0.0001 | 0.05 | Kruskal-Wallis; Dunn's correction |
| 6-S2C | WT vs RIM gap junction KD | ns | 0.5263 | 0.05 | Kruskal-Wallis; Dunn's correction |
| 6-S2C | WT vs *tdc-1*; RIM gap junction KD | ** | 0.0025 | 0.05 | Kruskal-Wallis; Dunn's correction |
| 6-S2D | WT vs *tdc-1* | *** | <0.0001 | 0.05 | 2-sample Kolmogorov-Smirnov; Bonferroni correction |
| 6-S2D | WT vs RIM gap junction KD | *** | <0.0001 | 0.05 | 2-sample Kolmogorov-Smirnov; Bonferroni correction |
| 6-S2D | WT vs *tdc-1*; RIM gap junction KD | *** | <0.0001 | 0.05 | 2-sample Kolmogorov-Smirnov; Bonferroni correction |
| 6-S2E | WT vs *tdc-1* | *** | <0.0001 | 0.05 | 2-sample Kolmogorov-Smirnov; Bonferroni correction |
| 6-S2E | WT vs RIM gap junction KD | *** | <0.0001 | 0.05 | 2-sample Kolmogorov-Smirnov; Bonferroni correction |
| 6-S2E | WT vs *tdc-1*; RIM gap junction KD | *** | <0.0001 | 0.05 | 2-sample Kolmogorov-Smirnov; Bonferroni correction |
| 6-S2F | WT vs *tdc-1* | *** | <0.0001 | 0.05 | 2-sample Kolmogorov-Smirnov; Bonferroni correction |
| 6-S2F | WT vs RIM gap junction KD | *** | <0.0001 | 0.05 | 2-sample Kolmogorov-Smirnov; Bonferroni correction |
| 6-S2F | WT vs *tdc-1*; RIM gap junction KD | *** | <0.0001 | 0.05 | 2-sample Kolmogorov-Smirnov; Bonferroni correction |
| 6-S2G | WT vs *tdc-1* | ns | 0.3591 | 0.05 | 2-sample Kolmogorov-Smirnov; Bonferroni correction |
| 6-S2G | WT vs RIM gap junction KD | *** | <0.0001 | 0.05 | 2-sample Kolmogorov-Smirnov; Bonferroni correction |
| 6-S2G | WT vs *tdc-1*; RIM gap junction KD | *** | <0.0001 | 0.05 | 2-sample Kolmogorov-Smirnov; Bonferroni correction |
| 6-S2H | WT vs *tdc-1* | *** | <0.0001 | 0.05 | 2-sample Kolmogorov-Smirnov; Bonferroni correction |
| 6-S2H | WT vs RIM gap junction KD | *** | <0.0001 | 0.05 | 2-sample Kolmogorov-Smirnov; Bonferroni correction |
| 6-S2H | WT vs *tdc-1*; RIM gap junction KD | *** | <0.0001 | 0.05 | 2-sample Kolmogorov-Smirnov; Bonferroni correction |
| 7B | RIM::HisCl: WT (- his) vs WT (+ his) | ns | 0.1707 | 0.05 | Kruskal-Wallis; Dunn's correction |
| 7B | RIM::HisCl: WT (- his) vs RIM GJ KD (- his) | * | 0.0420 | 0.05 | Kruskal-Wallis; Dunn's correction |
| 7B | RIM::HisCl: WT (- his) vs RIM GJ KD (+ his) | ns | >0.9999 | 0.05 | Kruskal-Wallis; Dunn's correction |
| 7C | RIM::HisCl: WT (- his) vs WT (+ his) | *** | 0.0002 | 0.05 | 2-sample Kolmogorov-Smirnov; Bonferroni correction |
| 7C | RIM::HisCl: WT (- his) vs RIM GJ KD (- his) | *** | <0.0001 | 0.05 | 2-sample Kolmogorov-Smirnov; Bonferroni correction |
| 7C | RIM::HisCl: WT (- his) vs RIM GJ KD (+ his) | ns | 0.9579 | 0.05 | 2-sample Kolmogorov-Smirnov; Bonferroni correction |
| 7E | RIM::ReaChR: WT vs RIM glu KO | ** | 0.0022 | 0.05 | 2-sample Kolmogorov-Smirnov; Bonferroni correction |
| 7E | RIM::ReaChR: WT vs *tdc-1* | *** | 0.0002 | 0.05 | 2-sample Kolmogorov-Smirnov; Bonferroni correction |
| 7E | RIM::ReaChR: WT vs RIM glu KO; *tdc-1* | *** | <0.0001 | 0.05 | 2-sample Kolmogorov-Smirnov; Bonferroni correction |
| 7G | RIM::ReaChR: WT vs RIM gap junction KD | *** | <0.0001 | 0.05 | 2-sample Kolmogorov-Smirnov |
| 7-S1B; left | RIM::HisCl: WT (- his) vs WT (+ his) | *** | <0.0001 | 0.05 | 2-sample Kolmogorov-Smirnov; Bonferroni correction |
| 7-S1B; left | RIM::HisCl: WT (- his) vs *tdc-1* (- his) | *** | <0.0001 | 0.05 | 2-sample Kolmogorov-Smirnov; Bonferroni correction |
| 7-S1B; left | RIM::HisCl: WT (- his) vs *tdc-1* (+ his) | *** | <0.0001 | 0.05 | 2-sample Kolmogorov-Smirnov; Bonferroni correction |
| 7-S1B; right | RIM::HisCl: WT (- his) vs WT (+ his) | ** | 0.0026 | 0.05 | 2-sample Kolmogorov-Smirnov; Bonferroni correction |
| 7-S1B; right | RIM::HisCl: WT (- his) vs *tdc-1* (- his) | ns | 0.2085 | 0.05 | 2-sample Kolmogorov-Smirnov; Bonferroni correction |
| 7-S1B; right | RIM::HisCl: WT (- his) vs *tdc-1* (+ his) | ns | 0.1635 | 0.05 | 2-sample Kolmogorov-Smirnov; Bonferroni correction |
| 7-S1D; left | RIM::HisCl: WT (- his) vs WT (+ his) | ** | 0.0015 | 0.05 | 2-sample Kolmogorov-Smirnov; Bonferroni correction |
| 7-S1D; left | RIM::HisCl: WT (- his) vs RIM GJ KD (- his) | ns | 0.0897 | 0.05 | 2-sample Kolmogorov-Smirnov; Bonferroni correction |
| 7-S1D; left | RIM::HisCl: WT (- his) vs RIM GJ KD (+ his) | *** | 0.0001 | 0.05 | 2-sample Kolmogorov-Smirnov; Bonferroni correction |
| 7-S1D; right | RIM::HisCl: WT (- his) vs WT (+ his) | * | 0.0102 | 0.05 | 2-sample Kolmogorov-Smirnov; Bonferroni correction |
| 7-S1D; right | RIM::HisCl: WT (- his) vs RIM GJ KD (- his) | ns | 0.1463 | 0.05 | 2-sample Kolmogorov-Smirnov; Bonferroni correction |
| 7-S1D; right | RIM::HisCl: WT (- his) vs RIM GJ KD (+ his) | ns | >0.9999 | 0.05 | 2-sample Kolmogorov-Smirnov; Bonferroni correction |
| 7-S2A; left | RIM::ReaChR: WT vs RIM glu KO | ns | >0.9999 | 0.05 | Kruskal-Wallis; Dunn's correction |
| 7-S2A; left | RIM::ReaChR: WT vs *tdc-1* | ns | >0.9999 | 0.05 | Kruskal-Wallis; Dunn's correction |
| 7-S2A; left | RIM::ReaChR: WT vs RIM glu KO; *tdc-1* | ns | >0.9999 | 0.05 | Kruskal-Wallis; Dunn's correction |
| 7-S2A; right | RIM::ReaChR: WT vs RIM glu KO | ns | 0.7543 | 0.05 | Kruskal-Wallis; Dunn's correction |
| 7-S2A; right | RIM::ReaChR: WT vs *tdc-1* | ns | >0.9999 | 0.05 | Kruskal-Wallis; Dunn's correction |
| 7-S2A; right | RIM::ReaChR: WT vs RIM glu KO; *tdc-1* | ns | 0.7671 | 0.05 | Kruskal-Wallis; Dunn's correction |
| 7-S2B | RIM::ReaChR: WT vs RIM glu KO | *** | <0.0001 | 0.05 | 2-sample Kolmogorov-Smirnov; Bonferroni correction |
| 7-S2B | RIM::ReaChR: WT vs *tdc-1* | *** | <0.0001 | 0.05 | 2-sample Kolmogorov-Smirnov; Bonferroni correction |
| 7-S2B | RIM::ReaChR: WT vs RIM glu KO; *tdc-1* | *** | <0.0001 | 0.05 | 2-sample Kolmogorov-Smirnov; Bonferroni correction |
| 7-S2C | RIM::ReaChR: WT vs RIM glu KO | ns | 0.1260 | 0.05 | 2-sample Kolmogorov-Smirnov; Bonferroni correction |
| 7-S2C | RIM::ReaChR: WT vs *tdc-1* | ns | 0.5967 | 0.05 | 2-sample Kolmogorov-Smirnov; Bonferroni correction |
| 7-S2C | RIM::ReaChR: WT vs RIM glu KO; *tdc-1* | ns | 0.0501 | 0.05 | 2-sample Kolmogorov-Smirnov; Bonferroni correction |
| 7-S2D; left | RIM::ReaChR: WT vs RIM gap junction KD | ns | 0.5512 | 0.05 | Mann-Whitney |
| 7-S2D; right | RIM::ReaChR: WT vs RIM gap junction KD | *** | 0.0005 | 0.05 | Mann-Whitney |
| 7-S2E | RIM::ReaChR: WT vs RIM gap junction KD | *** | <0.0001 | 0.05 | 2-sample Kolmogorov-Smirnov |
| 7-S2F | RIM::ReaChR: WT vs RIM gap junction KD | *** | 0.0002 | 0.05 | 2-sample Kolmogorov-Smirnov |
